# Chromatin Topology Reorganization and Transcription Repression by PML/RARα in Acute Promyeloid Leukemia

**DOI:** 10.1101/2020.03.05.979070

**Authors:** Ping Wang, Zhonghui Tang, Byoungkoo Lee, Jacqueline Jufen Zhu, Liuyang Cai, Przemyslaw Szalaj, Simon Zhongyuan Tian, Meizhen Zheng, Dariusz Plewczynski, Xiaoan Ruan, Edison T. Liu, Chia-Lin Wei, Yijun Ruan

## Abstract

**Background:** Acute promyeloid leukemia (APL) is characterized by the oncogenic fusion protein PML/RARα, a major etiological agent in APL. However, the molecular mechanisms underlying the role of PML/RARα in leukemogenesis remains largely unknown.

**Results:** Using an inducible system, we comprehensively analyzed the 3D genome organization in myeloid cells and its reorganization after PML/RARα induction, and performed additional analyses in patient-derived APL cells with native PML/RARα. We discovered that PML/RARα mediates extensive chromatin interactions genome-wide. Globally, it redefines the chromatin topology of the myeloid genome toward a more condensed configuration in APL cells; locally, it intrudes RNAPII-associated interaction domains, interrupts myeloid-specific transcription factors binding at enhancers and super-enhancers, and leads to transcriptional repression of genes critical for myeloid differentiation and maturation.

**Conclusions:** Our results not only provide novel topological insights for the roles of PML/RARα in transforming myeloid cells into leukemia cells, but further uncover a topological framework of a molecular mechanism for oncogenic fusion proteins in cancers.

## Introduction

Leukemias are often triggered by chromosomal rearrangements, such as translocations and inversions, which can generate oncogenic fusion transcription factors [1,2]. A hallmark in acute promyeloid leukemia (APL) is a chromosomal translocation that fuses the promyelocytic leukemia gene (*PML*) on chromosome 15 and the retinoic acid receptor alpha gene (*RARα*) on chromosome 17 into a fusion gene *PML/RARα* [3,4]. This translocation, denoted as t(15;17)(q24;q21), occurs in 98% of APL patients, and this fusion gene encodes a fusion protein PML/RARα, considered a major etiological agent of APL. In normal myeloid cells, RARα (a nuclear receptor and transcription factor) plays important roles in myelopoiesis, especially in granulocytic and monocytic differentiation programs [5,6]. However, the fusion protein PML/RARα in APL has been suggested to compete with endogenous RARα for binding at the same RA response elements (RAREs), which in turn leads to repression of normal RARα signaling in a dominant negative manner [7]. It has also been hinted that PML/RARα could predominantly target promoters regulated by transcription factor PU.1 through protein-protein interactions between PU.1 and RAREh binding sites genome-wide [8]. Early studies have suggested that PML/RARα may also abnormally recruit a histone deacetylase (HDAC) and/or polycomb repressive complexes (PRCs) to target genes important in hematopoietic differentiation [9,10], indicating that PML/RARα may have a role in altering chromosome configuration during APL genesis. Taken together, these investigations suggest a significant range of genome-wide restructuring induced by PML-RARα; however, how the comprehensive molecular mechanisms underlying the role of PML/RARα in leukemogenesis remain largely unknown.

Over the last decade, it has become clear that the human genomes are folded in complex 3-dimensional (3D) organizations in nuclei, and that 3D chromatin architectrures may be important in the higher order regulation of transcription regulation [11]. Several studies have hinted that chromosomal rearrangements in acute myeloid leukemia (AML) with inv(3)/t(3,3) lead to long-range interactions characterized by the 3D repositioning of a *GATA2* enhancer to the *EVI1* promoter to ectopically activate *EVI1*, which can cause dysregulation of both genes, with AML as the outcome [12,13]. Another study has shown that the deletion of insulated chromatin domain boundaries could activate proto-oncogene expression through aberrant distal regulatory elements, thereby contributing to T-cell acute lymphoblastic leukemia (T-ALL) [14]. Although these reports togenter indicated that chromatin configuration change might be an important feature in the transformation of normal cells into leukemic cells by oncogenic fusion proteins, specific evidence is lacking.

Considering that RARα binds directly to DNA genome-wide, we posit that the oncogenic fusion protein PML/RARα may also possess chromatin interaction properties, and alter the 3D genome topology as an critical event during leukemogenesis. To this end, we comprehensively analyzed normal myeloid cells with inducible PML/RARα, and patient-derived APL cells with native PML/RARα, to determine the roles of PML/RARα in 3D genome organization and transcription regulation, using integrative approaches including ChIA-PET for chromatin interactions, ChIP-seq for epigenomic states, and RNA-seq for transcriptional outputs. We discovered that PML/RARα mediated extensive long-range chromatin interactions genome-wide, distorted the established chromosomal folding topology in normal myeloid cells, and specifically repressed transcription of genes that are important to myeloid differentiation and maturation, together suggesting a topological mechanism for PML/RARα in leukemogenesis.

## Results

### Genome-wide chromatin interactions in normal myeloid cells and APL cells

To investigate the mechanisms through which PML/RARα leads to the development of promyelocytic leukemia phenotypes in myeloid cells, we employed a well-established PR9 cell line, which is dereived from U937 myeloid precursors at the promonocytic stage. PR9 cell line possesses normal endogenous PML and RARα genes, but also contains a transgenic construct inducible for PML/RARα expression via addition of ZnSO4. Upon induction, the expression of PML/RARα in PR9 drives the cells to develop promyelocytic leukemia phenotypes [15]. It has also been shown that the protein expression level of PML/RARα in ZnSO4-treated PR9 cells is comparable to that in APL patient-derived NB4 cells [8]. Hence, by comparing PR9 cells under normal condition (without PML/RARα protein) with PR9 cells under ZnSO4-induction conditions (with induced PML/RARα), we can investigate the dynamic changes of the myeloid genome mediated by the nacent fusion protein PML/RARα in PR9 cells (Fig. 1a).

**Figure 1.**
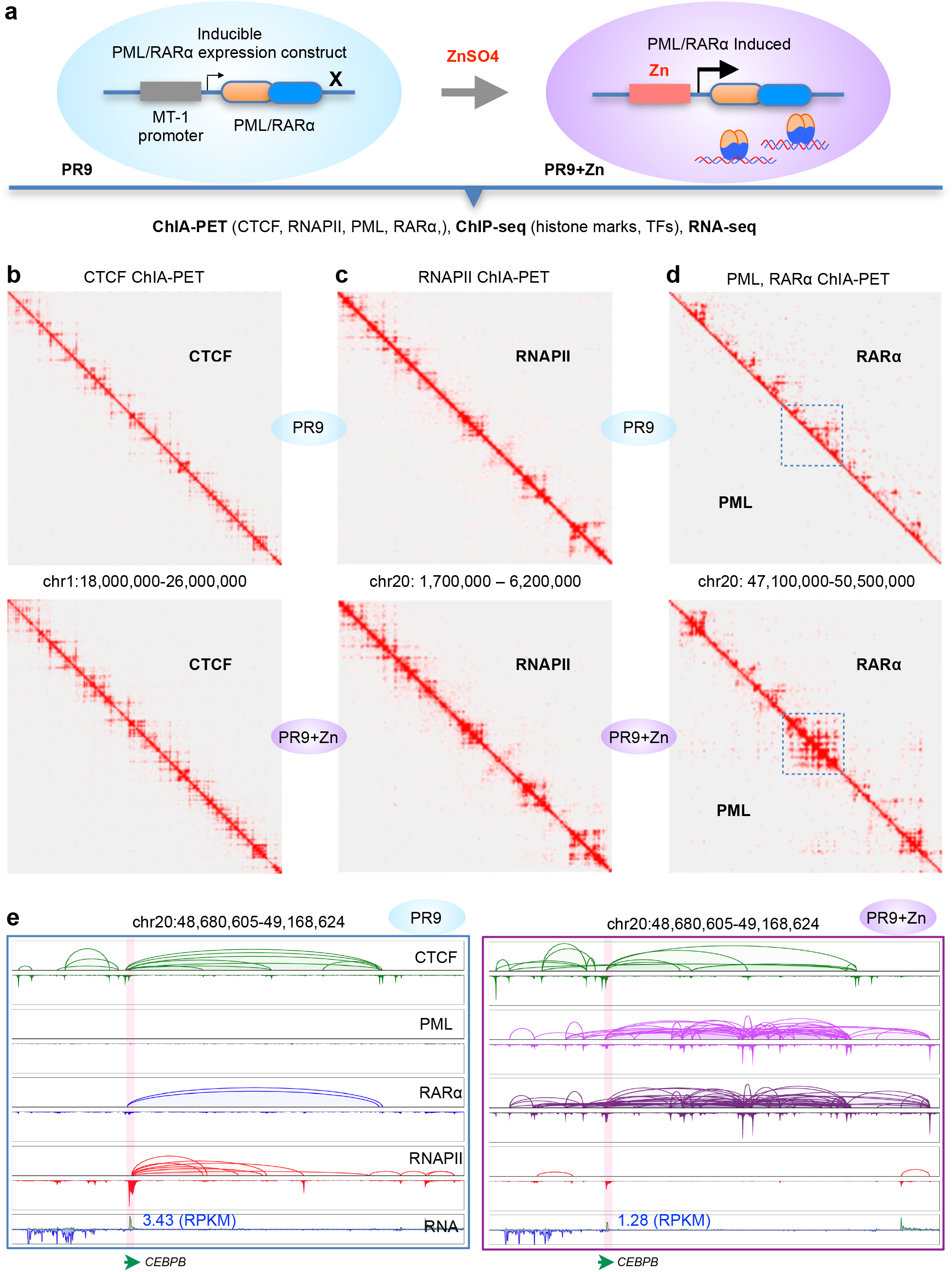
Mapping of 3D epigenome organizations in myeloid cells with inducible PML/RARα. a. Schematic of experimental designs using an inducible system. PR9 cells contain an inducible construct of the fusion gene PML/RARα. Upon ZnSO4 induction, the fusion gene will be activated, and the PML/RARα protein is expressed and interacts with the myeloid genome. Both PR9 and PR9+Zn cells were analyzed by ChIA-PET, ChIP-seq, and RNA-seq to map the 3D epigenomes. b. 2D chromatin contact maps of the PML and RARα ChIA-PET data from PR9 (top) and PR9+Zn (bottom) cells. To be noted, PML ChIA-PET did not produce meaningful data, while the RARα ChIA-PET generated abundant data mapping RARα-mediated chromatin interactions in PR9 cells (top). However, both PML and RARα experiments in PR9+Zn cells produced equal amounts of data with same patterns, indicating the detection of chromatin interactions mediated by the fusion protein PML/RARα. The boxed segments were zoomed-in for details in E. c. 2D chromatin contact maps of the CTCF ChIA-PET data from PR9 (top) and PR9+Zn (bottom) cells. d. 2D chromatin contact maps of the RNAPII ChIA-PET data from PR9 (top) and PR9+Zn (bottom) cells. e. Screenshots of browser views displaying a genomic segment at *CEBPB* loci in chr20, exemplifying the chromatin interactions detected by ChIA-PET using PML, RARα, CTCF, and RNAPII antibodies in PR9 (left) and PR9+Zn (right) cells. As shown, the CTCF data (green) exhibited same patterns and intensity in both PR9 and PR9+Zn cells; No PML data (pink) in PR9 cells, but extensive data in PR9+Zn cells; weak RARα signals in PR9 but strong strong and abundant (purple) in PR9+Zn; strong RNAPII (red) binding at the CEBPB gene locus and and interaction loops to enhancer sites in PR9 cells, but absent in PR9+Zn cells. The RNA-seq data showing that CEBPB is expressed in PR9 cells, but reduced by twofold in PR9+Zn cells.

To map the PML/RARα-initiated chromatin interactions to the myeloid genome, we performed ChIA-PET experiments using anti-PML and anti-RARα antibodies in both PR9 and PR9+Zn cells (Supplementary Table 1). Although the PR9 cells carry a transgenic PML/RARα construct, they would not express PML/RARα fusion protein without ZnSO4 induction, and should retain the endogenous *PML* and *RARα* genes and normal expression of native PML and RARα proteins. After ZnSO4 treatment, hereafter referred to as PR9+Zn cells, the PR9+Zn cells should acquire the induced PML/RARα protein while still retain the native PML and RARα proteins. Therefore, the PML- and RARα-enriched ChIA-PET experiments were expected to detect both of the native proteins and the induced PML/RARα in PR9+Zn cells, but only the native proteins in PR9 ocells. As shown in the 2D chromatin contact profiles of the ChIA-PET data (Fig. 1b), the RARα ChIA-PET experiment generated distinctive and typical chromatin contact data that mapped along the 2D contact diagonal, whereas the PML ChIA-PET experiment produced no meaningful chromatin contact data. This was expected, as RARα is a DNA-binding transcription factor, and PML does not interact with chromatin in nuclei. In contrast, we obtained extensive chromatin contact data in both the PML and RARα ChIA-PET experiments, from PR9+Zn cells (Fig. 1b). The striking similarity of chromatin contact patterns exhibited by these two ChIA-PET experiments in PR9+Zn cells using different antibodies (anti-PML and anti-RARα) validates that the induced fusion protein PML/RARα mediated new chromatin interactions in PR9+Zn cells.

To systematically investigate the impact of PML/RARα on the genomes, it is neccessary to characterize the 3D genome organization in both normal myeloid cells and APL cells. Therefore, we first generated high-quality CCCTC-binding factor (CTCF)-enriched ChIA-PET data (Supplementary Table 1) and mapped the higher-order chromosomal folding architectures and the detailed chromatin domain topology mediated by CTCF in PR9 cells and PR9 cells with induced PML/RARα expression *via* addition of ZnS04. Overall, these two CTCF datasets were highly correlated (Supplementary Fig. 1a-b), and the 2D contact profiles appeared to be identical (Fig. 1c), indicating that the ZnSO4 treatment did not directly alter the CTCF-mediated chromatin interactions in the myeloid genome.

In addition, we also performed RNA polymerase II (RNAPII) ChIA-PET experiments (Table S1) to map transcriptional chromatin interactions involving promoters and enhancers for active genes in PR9 and PR9+Zn cells. Globally, the two RNAPII ChIA-PET datasets appeared highly comparable (Fig. 1d; Supplementary Fig. 1c-d); however, locally, we observed significant difference (Fig. 1e). For example, at the locus of the gene *CEBPB* (encoding an important transcription factor for myeloid differentiation), it is observed that although the CTCF-mediated chromatin interactions in PR9 and PR9+Zn cells were very similar, the overwhelming chromatin contacts mediated by PML/RARα (detected by PML and RARα ChIA-PET data) in PR9+Zn cells appeared to overwrite the normal chromatin foliding architecture pre-defined by CTCF around the *CEBPB* (Fig. 1e). Remarkably, the extensive RNAPII occupancy at the *CEBPB* promoter and the abundant looping contacts from the promoter to enhancers shown in PR9 cells (Fig. 1e left) were abolished in PR9+Zn cells (Fig. 1e right). Consequently, the transcription of *CEBPB* was repressed in PR9+Zn cells as measured by RNA-seq data. Together, these observations imply that PML/RARα could potentially have a strong impact on the chromatin folding architecture and transcription regulation in myeloid cells.

### Topological reorganization of the myeloid genome by PML/RARα

To meticulously characterize the myeloid 3D genome organization and the impact of PML/RARα on myeloid genome topology, we first comprehensively characterized the CTCF ChIA-PET data in PR9 and PR9+Zn cells for their 3D chromatin organization, which reflected the native genome status of the myeloid genome. Because the two CTCF datasets were highly consistent (Fig. 1c) and that the ZnSO4 treatment in PR9 cells did not significantly change the CTCF chromatin interaction domains in myeloid genome, we combined the two CTCF ChIA-PET datasets for increased data coverage to create a reference topological map mediated by CTCF of the myeloid genome in PR9 cells. Based on the connectivity of the CTCF loop clusters, we identified 2,699 CTCF contact domains (CCD) covering the majority of the myeloid genome (Fig. 2a), which is comparable with the CCDs previously detected in the genome of B-lymphoblastoid GM12878 cells [16].

**Figure 2.**
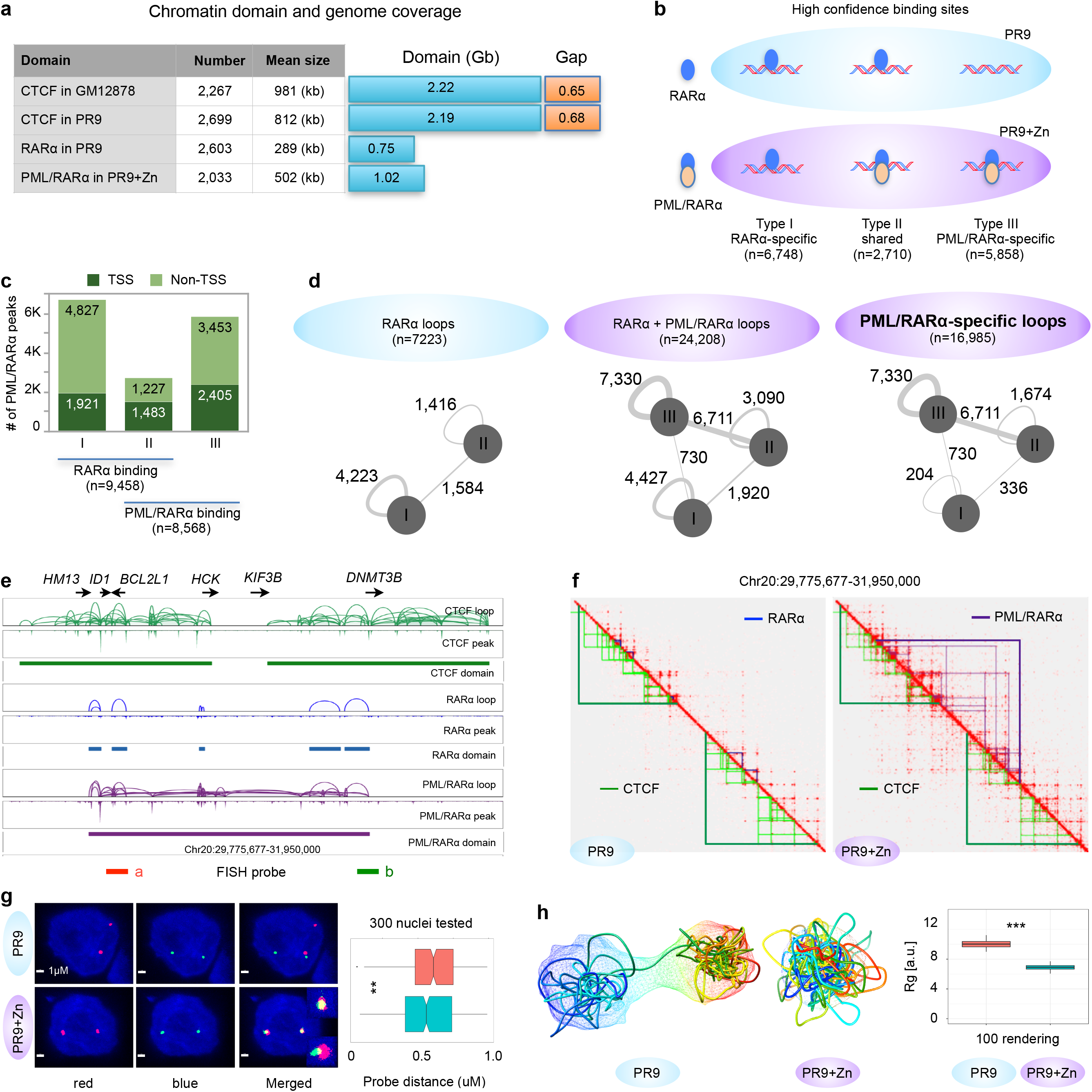
Reorganization of chromatin topology by PML/RARα-mediated chromatin interactions. a. Chromatin domains and genomic coverages by CTCF (combined data in PR9 and PR9+Zn cells), RARα (PR9) and PML/RARα (combined data in PR9+Zn). Summary table shows domain numbers and size of interaction domains. CTCF data from B-lymphoblastoid cells (GM12878) was given as a reference. b. Schematic of RARα and PML/RARα binding sites detected in PR9 and PR9+Zn cells detected by RARα- and PML-ChIA-PET experiments. RARα-specific (I), RARα and PML shared (II), and PML/RARα-specific binding sites were identified based on the combination patterns of the data in both PR9 and PR9+Zn cells. c. Characterized RARα and PML/RARα binding sites in relation to TSS of genes. d. Identification and classificiation of RARα- and PML/RARα-associated chromatin interactions in PR9 and PR9+Zn cells. Three types of RARα and PML/RARα binding sites (b) involved in interactions are indicated as circles. The interactions between two sites are indicated with lines. The thickness of lines corresponds to the number of interactions. Numbers of chromatin interactions in each category are given alongside the lines. e. Screenshot of genome browser view of chromatin interaction loop, binding peak and chromatin interaction domain for CTCF (green), RARα (blue), and PML/RARα (purple) at the *ID1-HCK* locus. Binding and looping signals for each protein factor were normalized. f. Integrated 2D chromatin contact maps for genomic segment (same as in E) on chr20 for CTCF, RARα, and PML/RARα ChIA-PET data from PR9 and PR9+Zn cells. The red signals in the contact maps were from the combined CTCF and RARα data (left, PR9 cells) and the combined CTCF, RARα, and PML data (right, PR9+Zn cells). Light green and dark green triangles depict CTCF loop and CCD, respectively; light blue and dark blue triangles depict RARα loops and domains; light and dark purple depict PML/RARα loops and domains. Red triangles indicate RNAPII-mediated loops and domains. g. 3D DNA-FISH validation. Two probes (red and blue) were designed at the corresponding position in the two separated CTCF domains as shown in E. Left panel: Example 3D DNA-FISH images of separated and merged views for the two probes (A in red, B in green) were shown in both PR9 and PR9+Zn cells. Right panel: Boxplot of spatial distance between the two probes measured microscopically from 300 nuclei in each of the PR9 and PR9+Zn cells. Mann-Whitney u test was used to test difference. ** *p* < 0.01. h. 3D chromatin folding rendering. Simulated 3D models of average structure and ensemble cloud in PR9 (left) and PR9+Zn (middle) using the data in corresponding region in F. Boxplot of radial diameter of simulated 3D models. K-S test was used to test differences. ** *p* < 2.2e-16.

We then analyzed the RARα ChIA-PET and PML ChIA-PET data in PR9 and PR9+Zn cells. The analysis of the RARα ChIA-PET data In PR9 cells to detect endogenous RARα is straightforward, same as we did for CTCF and RNAPII ChIA-PET data. However, in PR9+Zn cells, it is complicated, because both of the native RARα and the induced PML/RARα were expressed. Therefore, to distinguish the PML/RARα-associated chromatin contacts from the RARα-associated contacts, we dissected the ChIA-PET data (RARα) from PR9 cells and the data (PML and RARα) from PR9+Zn cells based on binding sites and chromatin loops. Hereby, we first analyzed the protein binding sites of the ChIA-PET data in both PR9 (with native RARα) and PR9+Zn cells (with native RARα and induced PML/RARα). To filter for high-confidence data, we defined a reliable binding site that was supported by at least two of the three independent ChIA-PET datasets (RARα data in PR9, RARα in PR9+Zn, and PML in PR9+Zn). Using this criterion, we identified 9,458 RARα binding sites in PR9 cells and 8,568 PML/RARα binding loci in PR9+Zn cells (Fig. 2b). The majority of the RARα binding sites (6,748; 71%) were RARα-specific (detected in both PR9 and PR9+Zn cells, but not in PML data in PR9+Zn cells). However, there were 2,710 (29%) RARα binding sites in PR9 cells that were also found in PML and RARα data in PR9+Zn cells, suggesting a possible co-occupancy or a competiton mode at those loci by the native RARα and the induced PML/RARα. In addition, we identified 5,858 PML/RARα-specific loci in PR9+Zn cells (Fig. 2b). Overall, both the native RARα and fusion protein PML/RARα demonstrated similar genome-wide chromatin binding capacity (9,458 vs. 8,568). Proportionally, 28% (1,921 / 6,748) of the RARα-specific binding sites were located proximal to gene transcription start sites (TSS), whereas 41% (2,405 / 5,858) of the PML/RARα-specific binding sites and 55% (1,483 / 2,719) of PML/RARα sites co-localized with RARα sites were proximal to TSS (Fig. 2c). This observation suggests an increased tendency for PML/RARα to target to gene promoters for alteration of gene transcription regulation, in addition to binding at non-genic regions to impact chromatin architecture.

Next, we analyzed the chromatin contact loops mediated by RARα or PML/RARα, which identified 7,223 high-confidence loops by RARα in PR9 cells, and 24,208 loops in PR9+Zn cells by a combinatorial effects of native RARα and the induced nascent PML/RARα (Fig. 2d). Intriguingly, although the binding capacities of RARα and PML/RARα were similar (Fig. 2b), extensive chromatin interactions detected in PR9+Zn cells were largely associated with PML/RARα binding loci, particularly the PML/RARα-specific binding loci. Collectively, our observations imply a substantial impact from PML/RARα to alter the topological architecture of the myeloid genome.

To investigate how PML/RARα affects the myeloid genome, we aggregated the PML/RARα-associated chromatin interactions into PML/RARα contact domains, similarly to what we did for CTCF domains (Fig. 2a). Intriguingly, when integrating the PML/RARα complex domains into the CTCF-defined topological framework of the myeloid genome (Supplementary Fig. 2a), we found that many (249) PML/RARα chromatin domains extended across the boundaries of two adjacent CCDs and connected parts of them, resembling “stitches” weaving multiple CCDs together. These “stitch” PML/RARα complex domains usually involved high levels of chromatin contacts (Supplementary Fig. 2b), and were prevalently spread across the entire genome (Supplementary Fig. 2c). The resulting “stitched CCD” by PML/RARα exhibited extended domain coverage (Supplementary Fig. 2d), and thus potentially had a global impact on the overall topological organization of the myeloid genome. Such impacts were particularly visible at the level of topological domains. For example, at a 2.17 Mb segment of chr20, the two adjacent but separated CCD with scattered RARα binding and looping in PR9 cells were brought together by a PML/RARα chromatin domain with extensive binding and looping as shown in PR9+Zn cells (Fig. 2e-f). This observation was further validated by a two-color DNA-FISH experiment, showing that the two separated CTCF domains in PR9 cells were in much closer contacts in PR9+Zn cells than in PR9 cells (Fig. 2g). An ensemble structure-based algorithm was applied to the chromatin interaction data derived from PR9 and PR9+Zn cells (Supplementary Methods), and elucidated the topological structural changes resulting from the action of the induced PML/RARα (Fig. 2h). Another example of the topological changes in PR9 cells before and after the ZnSO4 induction of PML/RARα was at a 3 Mb segment on chr18 (Supplementary Fig. 2e-g). Taken together, our data demonstrated that the fusion oncoprotein PML/RARα acts through extensive chromatin binding and looping genome-wide, and results in strong ectopic chromatin interactions that extend across the boundaries of CTCF-defined chromatin architectures in normal myeloid cells, thus leading to the topological reorganization of the myeloid genome into aberrant configurations in APL cells.

### Alteration of gene expression by PML/RARα

Subsequently, we systematically analyzed the RNAPII ChIA-PET data in PR9 and PR9+Zn cells in relation to PML/RARα-defined chromatin domains, and found that large numbers of RNAPII-associated chromatin interaction sites (proximal or distal to TSS) were co-occupied by PML/RARα in PR9+Zn cells (Supplementary Fig. 3a), indicating that PML/RARα may also directly interfere with the transcription programs in myeloid cells. We then quantified RNAPII occupancy at these sites in PR9 cells before and after ZnSO4 induction of PML/RARα, to assess the effects of PML/RARα on RNAPII. While most of the loci showed insignificant changes after 4 hours of ZnSO4 induction, we identified 871 (20%) loci that exhibited significant reduction of RNAPII binding intensity (Fig. 3a). As analysis controls, less than 10% of the non-PML/RARα sites, including the RARα binding sites, showed changes in RNAPII occupancy, presumably due to systems noise. Therefore, our observation suggests that PML/RARα might specifically target a subset of RNAPII interaction loci and induce functions that repress gene transcription.

**Figure 3.**
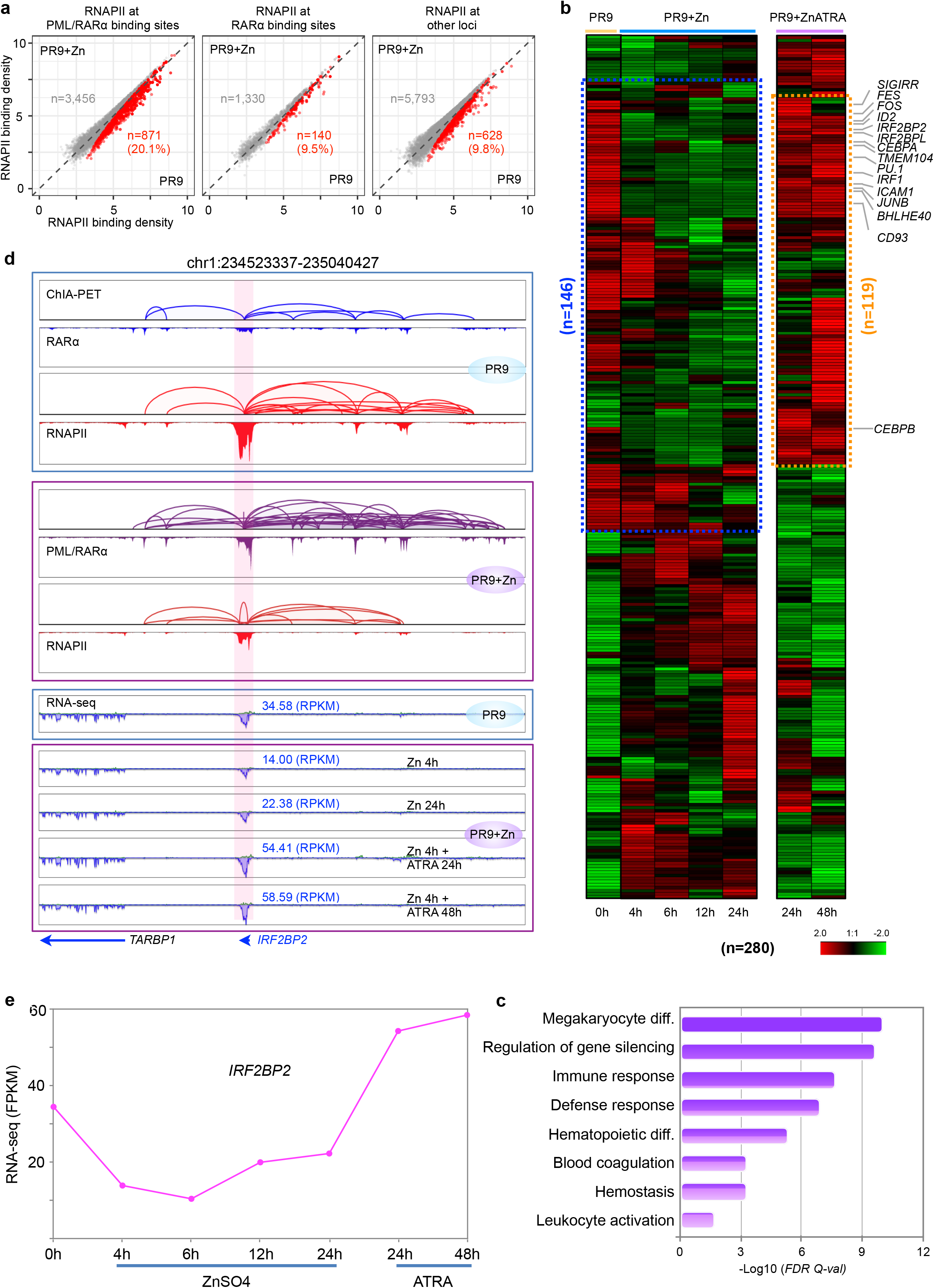
PML/RARα selectively disrupt RNAPII transcriptome. a. Scatter plots showing signal reduction of RNAPII occupancy at PML/RARα binding sites (left), RARα binding sites (middle), and other loci (right) in PR9+Zn cells. The red dots represent the data points of RNAPII binding intensity was significantly higher in PR9 than in PR9+Zn cells. The gray dots denote the RNAPII data points without significant changes in binding intensity between PR9 and PR9+Zn cells. The numbers of corresponding sites and the percentages of RNAPII changes are shown in each plot. b. Expression profiles of representative genes (n=280) whose promoters both exhibited decreased RNAPII binding intensity and overlapped with PML/RARα binding in PR9, PR9+Zn, and PR9+Zn+ATRA cells. The blue dashed box highlights the genes that were repressed by PML/RARα. The yellow dashed box highlights the genes (n=119) that were repressed by PML/RARα but rescued by ATRA treatment. Key genes known to be involved in leukemia biogenesis are indicated. c. Gene ontology (GO) enrichment analysis of genes (n=280) whose promoters represent decreased RNAPII binding intensity (control vs. treatment) in the PML/RARα category that were characterized in (A). X axis denotes enrichment score of −log10 (FDR). d. Screenshot of browser views of chromatin interaction loops and peaks for RARα (blue), RNAPII (red), and PML/RARα (purple) at the *SIGIRR* locus in PR9 and PR9+Zn cells. Strand-specific RNA-seq data in PR9 and PR9+Zn cells with time course of ATRA treatment. The *SIGIRR* region is highlighted. The expression data (RPKM) for *SIGIRR* expression are given in each RNA-seq track. e. Line plot shows mean expression level of *IRF2BP2* over the timepoints of ZnOS4 and ATRA treatments.

Next, we focused on the genes (n=288) that were associated with the reduced RNAPII occupancy due to co-occupancy by PML/RARα, and analyzed their transcription output using RNA-seq data over a timecourse during ZnSO4 induction of PML/RARα in PR9+Zn cells. Remarkably, more than half (n=146) of these genes exhibited a corresponding pattern of transcriptional reduction over the timecourse during ZnSO4 treatment (Fig. 3b). To test if the observed transcriptional repression was directly related to the induction of PML/RARα, we added all-trans retinoic acid (ATRA) to PR9+Zn cells in order to rescue the gene expression potentially hampered by the induced PML/RARα. ATRA is an important drug in APL treatment. It causes degradation of the PML/RARα fusion protein through the ubiquitin-proteosome and caspase system [17,18]. We therefore performed a “rescuing” experiment by adding ATRA to the PR9 cells that were under ZnSO4 induction of PML/RARα. Remarkably, in the “rescuing” experiments, most of the genes (81.5%; 119 / 146) were recovered by ATRA treatment, showing increased transcription (Fig. 3b; Supplementary Fig. 3b). Among these genes are many that are known for their functions involved in myeloid cell differentiation, including transcription factors and cytokines, such as the previously reported *CEBPB* [19], *ID2* [20], and *SPI1* [21] involved in megakaryocytic and granulocytic differentiation. Gene Ontology analysis to this set of genes showed that they are significantly enriched in biological processes associated with hemopoiesis, immune processes, myeloid cell activation and differentiation (Fig. 3c), further validating that at least part of the functions of PML/RARα is to act via repressing the transcription of genes involved in myeloid cell differentiation during APL pathogenesis.

As mentioned in Fig. 1e, the abundant RNAPII bindings and looping at the *CEBPB* promoter and its enhancer sites observed in PR9 cells were repressed by the induced PML/RARα in PR9+Zn cells while the CTCF-mediated chromatin folding structures unchanged, exemplifying a profound repressive function to transcription by PML/RARα. Simiarly, at the *IRF2BP2* locus, there was modest RARα ChIA-PET data and strong RNAPII-associated chromatin interactions between the *IRF2BP2* promoter and multiple enhancers detected in PR9 cells. However, after induction by ZnSO4, robust PML/RARα binding peaks and chromatin loops appeared, which directly overlapped with the RARα and RNAPII associated chromatin sites as detected in PR9 cells. Coincidentally, the RNAPII signals were much reduced in PR9+Zn cells (Fig. 3d). Furthermore, the RNA-seq data at this region showed more than 2-fold reduction of *IRF2BP2* expression when PML/RARα was induced in PR9+Zn cells, and, strikingly, rebounded after the addition of ATRA (Fig. 3d-e). Together, the high degree of correlation between the repression by ZnSO4 induction for PML/RARα expression and the liberation by ATRA treatment for PML/RARα degradation convincingly suggest that this set of genes may be the direct targets of PML/RARα for transcriptional repression in APL cells. It may also suggest mechanistically that PML/RARα forcefully compress the chromatin topological structure around myeloid-specific transcriptional cassette through its extensive chromatin binding and looping, and thus limit the access for transcription machinery.

### Interference with transcription factor binding at enhancer sites by PML/RARα

We reasoned that the potential specificity of PML/RARα targeting to a subset of actively transcribed genes in myeloid cells might be through interference with specific transcription factors (TF). It is widely known that TFs can facilitate the physical chromatin contacts between promoters and distal regulatory elements by looping the intervening DNA between them [22,23]. Specifically, PU.1 (also known as SPI1), known as a lymphoid-specific transcription activator, has been suggested to be associated with PML/RARα (Wang et al., 2010). To identify specific TFs involved at the PML/RARα chromatin interaction sites, we performed TF motif analysis (Supplemental Methods), and identified seven protein factors that were significantly enriched at PML/RARα binding sites (Supplementary Fig. 4a), including three TFs-PU.1, CEBPB, and IRF1 (Fig. 4a) -that are the most relevant and specific TFs in myeloid cells [8]. To further characterize these TFs, we performed ChIP-seq experiments and generated genome-wide binding profiles for PU.1, CEBPB, and IRF1 in PR9 and PR9+Zn cells, along with ChIP-seq of H3K9K14ac and P300 for promoters and enhancers. Interestingly, more than half of the TF binding sites found in PR9 cells were no longer present, or the binding signal intensities were significantly reduced, after PML/RARα induction in PR9+Zn cells (Fig. 4b). It is noteworthy that the binding profiles of these TFs were highly correlated with transcriptionally active marks for promoters (H3K9K14ac) and enhancers (P300), and RNAPII-associated chromatin interactions, higher in PR9 control cells (PR9) but lower in the cells under ZnSO4 induction of PML/RARα (PR9+Zn). For example, at the *PU.1* locus, PML/RARα bound specifically at the PU.1 promoter site and interacted with a number of enhancers, as indicated by H3K9K14ac and P300 binding profiles. in particular, at the enhancer sites, the occupancy by the three TFs (PU.1, CEBPB, and IRF1) were notably abolished or reduced in intensity (Fig. 4c). Simultaneously, the binding peaks for H3K9K14ac and P300 at the enhancers were also either decreased in signal intensity or abolished. The same observations were also exemplified at the *CEBPB* (Fig. 4d), and *IRF1* loci (Supplementary Fig. 4b).

**Figure 4.**
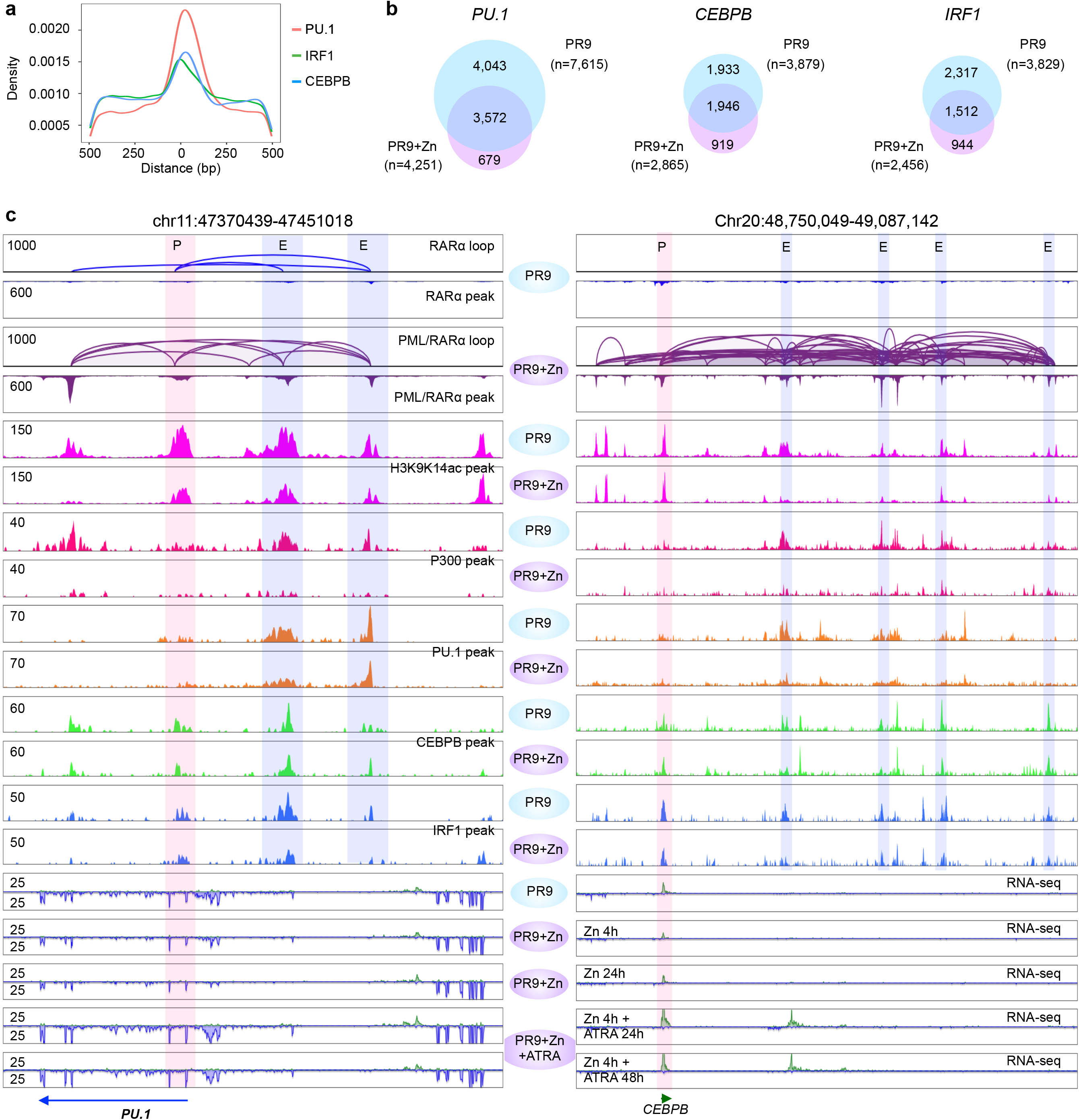
PML/RARα interrupted the transcription activity of key transcription factors. a. Motif enrichments of hematopoietic transcription factor at PML/RARα binding sites. b. Venn diagrams of the overlapped binding sites by three TFs (PU.1, CEBPB and IRF1) in PR9 and PR9+Zn cells. c. Two examples (left, on chr11; right, chr20) showing PML/RARα binding and looping, where the occupancy by H3K9K14ac (pink), P300 (red), PU.1 (orange), CEBPB (green), and IRF1 (blue) in PR9 cells was notably reduced in PR9+Zn cells. The promoter site is highlighted in light red and the enhancer sites are highlighted in light blue. RNA-seq data from PR9 cells and from the time course treatments in PR9+Zn cells are also shown.

Taken together, our results indicate that PML/RARα directly interacts with specific chromatin loci and disrupts the transcription of specific TFs. More importantly, the reduced TF activities likely further dysregulate the transcription programs of downstream target genes that are important for normal myeloid cell differentiation.

### Disruption of super enhancers by PML/RARα

The experiments above showing that PML/RARα disrupts the cobinding of multiple cofactors at RARα sites suggest that PML/RARα would perturb superenhancer function. Super-enhancers (SE), as a subset of regulatory elements, have been proposed to facilitate interactions between enhancers and promoters primarily associated with highly transcribed genes controlling cell identity and characteristically engaged multiple TFs at high intensity [24,25]. Previous studies have shown that SEs are critical in establishing and maintaining cell-specific transcriptional regulation of gene expression as well as fine-tuning of expression of many oncogenes [26,27]. Given that the induced PML/RARα prominently targeted at genomic regions with high levels of H3K9K14ac modification (Fig. 5a), we used the H3K9K14ac ChIP-seq data to catalogue SEs using the ROSE algorithm [26,27] in PR9 and PR9+Zn cells (Fig. 5b). In total, we identified 521 SEs overlapped with PML/RARα binding sites in PR9+Zn cells (Supplementary Fig. 5a), implying that PML/RARα may broadly interfere with the functions of SEs. Remarkably, of the 480 SEs identified in PR9 cells, more than half (n=247) lost their SE characteristics (H3K9K14ac signals) after Zn induction for PML/RARα PR9+Zn cells (Fig. 5c). It is observed that the RNAPII ChIA-PET data intensities (peaks and loops) associated with these SEs in PR9 cells were significantly decreased in PR9+Zn cells, in contrast, to the induction of PML/RARα and associated binding peaks and loops in PR9+Zn cells (Fig. 5d).

**Figure 5.**
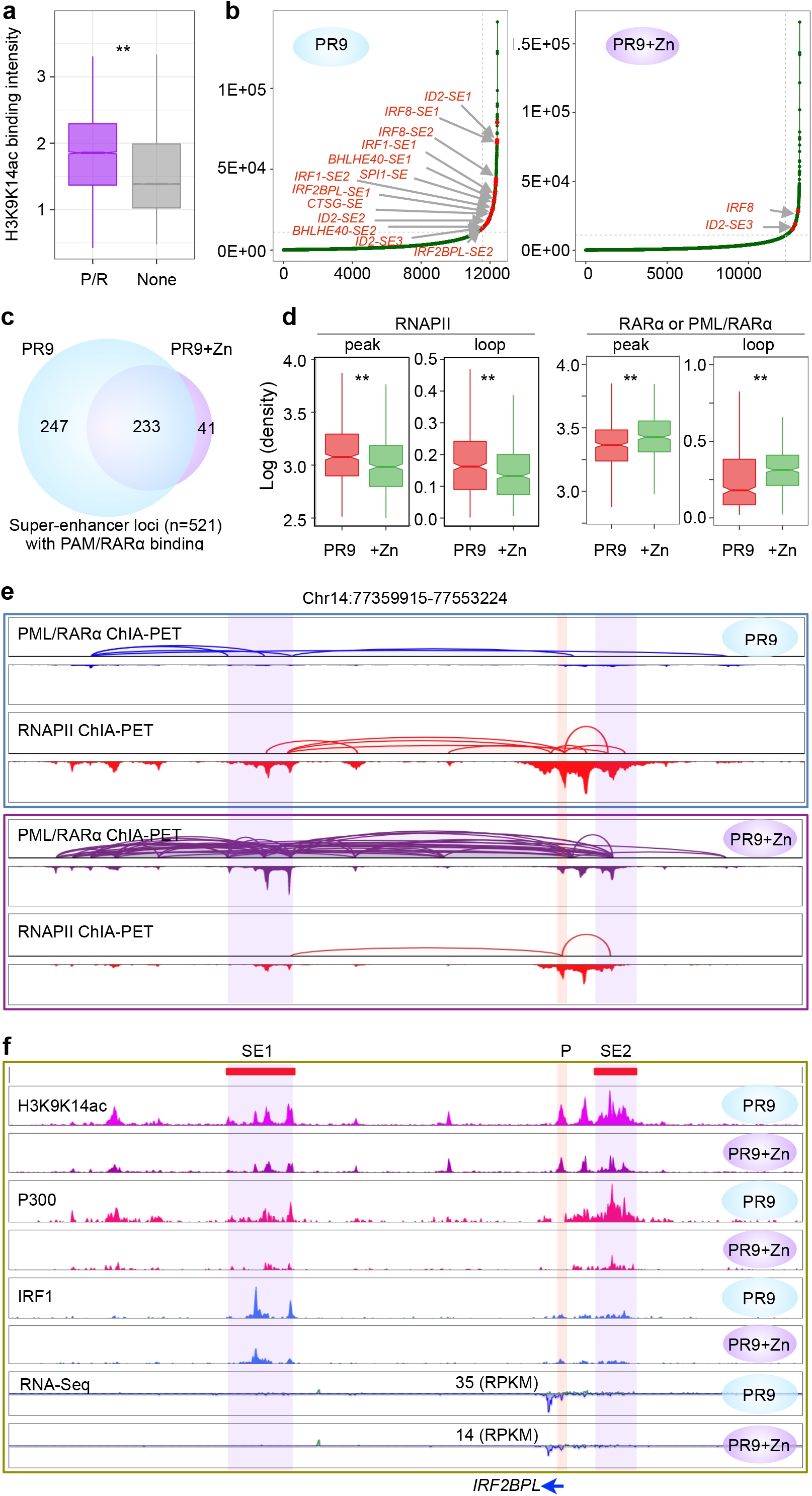
Super-enhancers affected by PML/RARα. a. Boxplots of H3K9K14ac ChIP-seq inntensity at the PML/RARα (P/R, purple) interaction binding sites or not as identified in PR9+Zn cells. The control (grey) represents H3K9K14ac signals at non-PML/RARα anchor loci (None). Paired t-test was used to test difference. ** p < 0.001. b. Distribution plots of H3K9K14ac ChIP-seq signals and the super-enhancers (SEs) identified in PR9 and PR9+Zn cells. SEs were ranked by increasing H3K9K14ac signals. SEs associated with genes critical in myeloid differentiation are highlighted in red. c. Venn diagram for the numbers of unique and common SEs in PR9 and ZnSO4-treated PR9 cells based on differential analysis of H3K9K14ac signals. d. Boxplots for normalized data signal intensity of RNAPII binding and loops at SE sites in PR9 and PR9+Zn (+Zn) cells (left), and the RARα binding and looping in PR9 cells and PML/RARα binding and looping in PR9+Zn (+Zn) cells at SE sites (right). ** *p* < 0.001. by Kolmogorov-Smirnov test. e. An example of chromatin interactions at the *IRF2BPL* locus identified by ChIA-PET of RARα (blue) and RNAPII (red) in PR9 cells, and PML/RARα (purple) and RNAPII (red) in PR9+Zn cells. Each ChIA-PET data are shown in tracks of loops (up) and peaks (below). f. At the same location as in E, two SEs (highlighted) were identified with clusters of multiple H3K9K14ac peaks in PR9 cells. The H3K9K14ac peak signals were notably reduced in PR9+Zn cells. Similarly, the ChIP-seq signals for P300 and IRF1obserevd in PR9 cells were aalso reduced in PR9+Zn cells. Also, the expression of *IRF2BPL* as measured by RNA-seq data (RPKM) in PR9 cells was reduced by more than twofold in PR9+Zn cells.

Based on the connectivity of RNAPII ChIA-PET data, we detected 282 genes that were linked to the PML/RARα-affected SEs (n=247). Subsequent GO analysis of this gene set identified 123 genes that were highly associated with functions in myeloid cell differentiation, and positive regulation of myeloid leukocyte differentiation and myeloid cell homeostasis (Supplementary Fig. 5b). Many of these 123 genes, including *FOS, IRF2BP2, ID2, IRF1, IRF2BPL*, and *BHLHE40*, were highly expressed in PR9 cells but repressed under ZnSO4 treatment conditions (Supplementary Fig. 5c), and as highlighted in Fig. 5b. In contrast, although 215 genes were found associated with the SEs (n=233) that were not affected by PML/RARα, most of those genes were not associated with myeloid specific functions (Supplementary Fig. 5b).

At the *IRF2BPL* locus, only base-level RARα signals but substantial RNAPII peaks and loops connecting the *IRF2BPL* promoter and enhancers in PR9 cells (Fig. 5e). However, following ZnSO4 induction, strong PML/RARα binding peaks and loops were observed in PR9+Zn cells overlapping directly with the RNAPII binding sites loops observed in PR9 cells. Consequently, the RNAPII signals in in PR9+Zn cells were significantly diminished compared to the RNAPII signals in PR9 cells (Fig. 5e). Evidently, the H3K9K14ac peak profile in this region called for two SEs (Fig. 5f) interconnecting with the *IRF2BPL* promoter by a substantial number of RNAPII loops in PR9 cells. Natably, the SE signals were substantially reduced in PR9+Zn cells, presumably by the PML/RARα effects. Similarly, P300, another enhacer mark showed the same pattern, with strong signals in PR9 cells, but diminished signals in PR9+Zn cells (Fig. 5f). Interestingly, the occupancy of IRF1 (a TF important for hematopoiesis) at the SE1 site was also much reduced in PR9+Zn cells. Correspondingly, the *IRF2BPL* expression was 2.5-fold down-regulated in PR9+Zn cells (Fig. 5f). Another example is at the *FOS* locus (Supplementary Fig. 5d).

Altogether, the above results demonstrated that PML/RARα may directly intrude super-enhancers, and the loss of properties of SEs may contribute to the disruption of RNAPII-mediated SE-to-promoters connectivity, consequently dysregulating gene transcription and alter the cell lineage controls during APL genesis.

### Native PML/RARα in patient-derived APL cells functions similarly to the inducible PML/RARα

To validate the above findings in the PR9 cellular system, we analyzed NB4 cells, a cell line derived from an APL patient harboring the t(15,17) translocation and expressing an native PML/RARα fusion protein [28]. Therefore, the cellular state of NB4 (with native PML/RARα) would be comparable to PR9+Zn cells (with induced PML/RARα). We also treated NB4 cells with ATRA (NB4+ATRA) to deplete the native PML/RARα. These NB4+ATRA cells (with PML/RARα depleted) thus match with the PR9 cells (no PML/RARα). We reasoned that, with these parallels between the NB4 and the PR9 inducible systems, a comparative analysis of the two systems would yield insights into the property and function differences between the inducible and native PML/RARα (Fig. 6a). First, we performed RNA-seq for gene expression analysis in the pairs of NB4 vs. PR9+Zn cells, and the NB4+ATRA vs. PR9 cells. The overall gene expression profiles between the two pairs exhibited high correlations (Fig. 6b, Supplementary Fig. 6a), indicating that the cellular systems of NB4 and PR9 were very comparable. However, when comparing the NB4 and NB4+ATRA cells, we observed that significant numbers of genes were up regulated, including many myeloid specific genes (Supplementary Fig. 6b). Sperifically to the set of genes (n=146) targeted by PML/RARα (Fig. 3b), all of them were up-regulated in NB4+ATRA cells (Fig. 6c), suggesting that native PML/RARα had similar effects to myeloid specific genes as in the PR9 cellualr system by the induced PML/RARα fusion protein.

**Figure 6.**
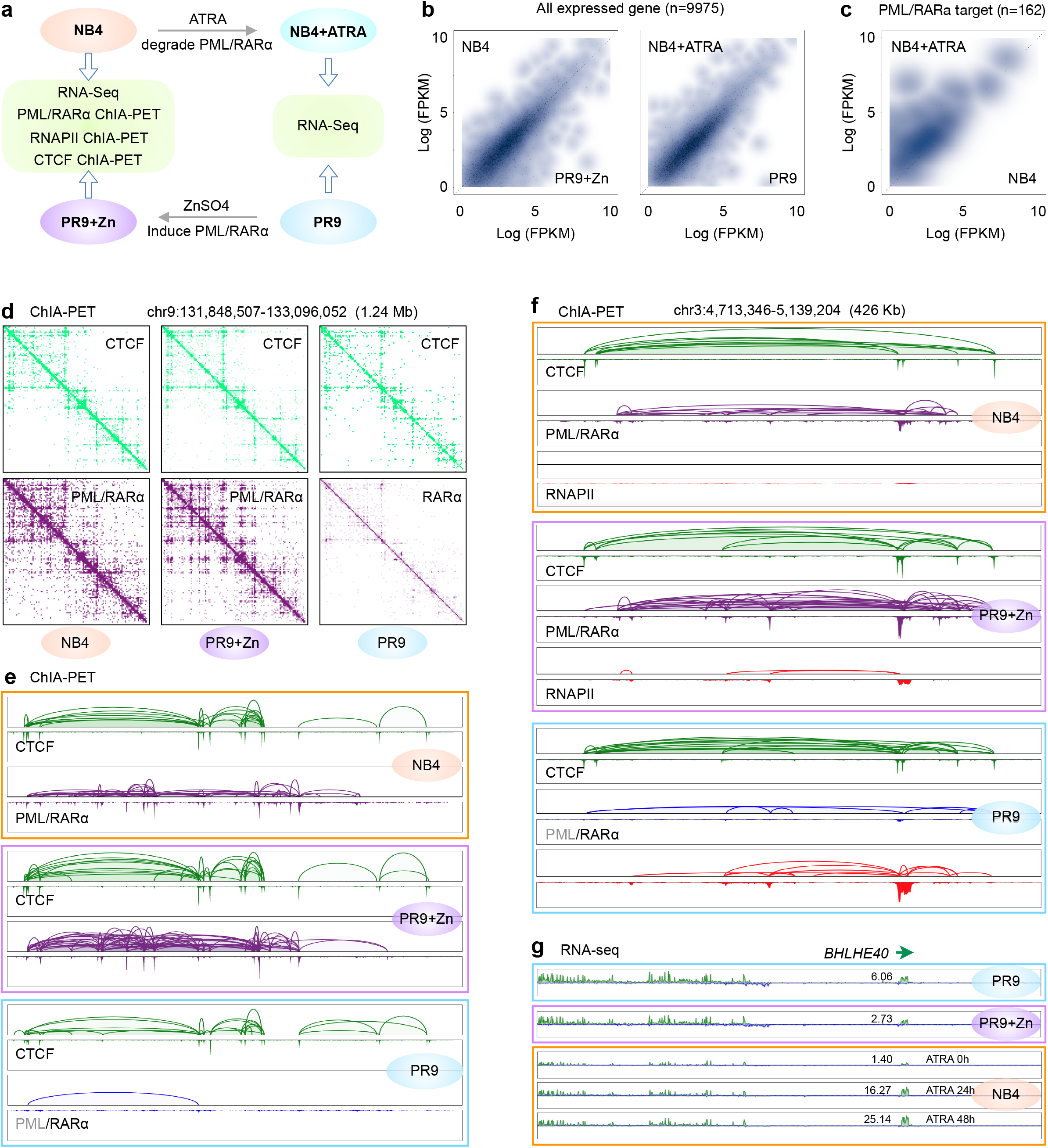
Native PML/RARα in patient-derived APL cells behaved the same as induced PML/RARα. a. Schematic design for comparison of transcription (RNA-seq) and 3D genome organization (ChIA-PET of PML/RARα, RNAPII, and CTCF) between patient-derived APL cells (NB4, with native PML/RARα) vs. PR9 cells by ZnSO4 induction (PR9+Zn, with induced PML/RARα), as well as for comparison of transcription (RNA-seq) between NB4 cells under ATRA treatment (NB4+ATRA, with native PML/RARα depleted) vs. PR9 cells without PML/RARα. b. Contour plots for correlation of gene expression (FPKM) between NB4 vs. PR9+Zn cells (left) and NB4+ATRA vs. PR9 cells (right). c. Contour plots for correlation of gene expression (FPKM) of the 146 PML/RARα target genes between NB4 vs. NB4+ATRA cells. d. Integrated 2D contact maps of PML/RARα (up, purple) and CTCF (low, green) ChIA-PET data in NB4 (left), PR9+Zn (middle), and PR9 (right) cells. e. A screenshot of browser view for chromatin loops and pleaks by CTCF (green) and PML/RARα (purple) showing high similarity between NB4 (top) and PR9+Zn (middle) cells. Chromatin loops and peaks by CTCF (green) and RARα (blue) in PR9 cells are included as a reference of normal myeloid cells. f. An example for comparison at the *BHLHE40* locus of chromatin loops peaks by CTCF (green), PML/RARα (purple) or RARα (blue), and RNAPII (red) in cells of NB4 (orange box), PR9+Zn (purple box), and PR9 (blue box) cells. It indicated that the chromatin structures mediated by CTCF, and PML/RARα, and RNAPII in NB4 cells and PR9+Zn cells were highly comparable. When compared to the data in PR9 cells, both NB4 and PR9+Zn cells exhibited high levels of PML/RARα chromatin interactions, but basal levels of RNAPII occupancy. g. At the same *BHLHE40* locus, the expression (FPKM) of gene *BHLHE40* as measured in PR9 were significantly reduced in PR9+ZN and NB4 cells, but recovered in NB4 cells afater ATRA treatments.

Next, we performed ChIA-PET analyses for protein factors CTCF, PML/RARα, and RNAPII in NB4 cells, and then compared with the same datasets derived from PR9+Zn (Fig. 6a; Fig. 1b-d). Similarly, the RNAPII and PML/RARα binding peak profiles between NB4 and PR9+Zn cells were also highly correlated (Supplementary Fig. 6b). We further analyszed the chromatin contacts of the ChIA-PET data. The 2D contact profiles of the PML/RARα ChIA-PET data obtained in NB4 cells appeared to be very similar to the PML/RARα data in PR9+Zn cells, and obviously different from the RARα (no PML) data in PR9 cells (Fig. 6d), clearly indicating that the native PML/RARα in NB4 cells behaved similarly to the induced PML/RARα in PR9+Zn cells. As a reference, the CTCF-mediated chromatin contacts in NB4, PR9+Zn and PR9 cells were highly comparable, as expected. More specifically, the CTCF loops and peaks were highly consistent in the three samples, and the PML/RARα loops and peaks were also consistent in NB4 and PR9+Zn cells, but not the same in PR9 cells, where there were no PML/RARα data except the data derived from RARα (Fig. 6e). Collectively, these observations suggested that the behavior of native PML/RARα in genome topological organization in NB4 cells was similar to that of the induced PML/RARα in PR9+Zn cells.

Furthermore, we observed that PML/RARα in NB4 cells also strongly inhibite RNAPII occupancy and transcriptional chromatin interactions at many myeloid-specific gene loci in PR9 cells (Supplementary Fig. 6a). For instance, at the *BHLHE40* locus in PR9 cells (Fig. 6f), RNAPII showed abundant occupancy at the gene promoter and mediated extensive chromatin loops to enhancers. However, at the same locus in PR9+Zn and NB4 cells, strong PML/RARα binding and looping were observed, and the RNAPII signals were diminished (Fig. 6f). To further investigate whether the native PML/RARα in NB4 cells affects the gene expression of myeloid-specific genes, as we showed in PR9 and PR9+Zn cells, we added ATRA to NB4 cells to deplete the native PML/RARα and then measured the transcripts by RNA-seq analysis. Differential expression analysis showed that many myeloid specific genes expressed at high levels in PR9 cells (*HCK, BHLHE40, CEBPB, IRF1*, etc) were repressed in NB4 cells and were reactivated after 24 hours and 48 hours of ATRA treatments (Supplementary Fig. 6a). For example, the normal expression of *BHLHE40* in PR9 cells was in a modest level (6.06 FPKM), and was repressed more than twofold (2.73 FPKM) after 4 hours of ZnSO4 induction of PML/RARα in PR9+Zn cells. At the same locus in NB4 cells, *BHLHE40* was repressed. However, after ATRA treatment, the expression of this gene increased more than tenfold (Fig. 6g). Another prominent example is at the *FOS* locus (Supplementary Fig. 6c). Together, these results further suggested that our observations for PML/RARα in the PR9 inducible system faithfully reflected the native PML/RARα functions for chromosomal reorganization and transcriptional repression in patient-derived APL cells.

## Discussion

In this study, we comprehensively mapped the 3D geome organizations and epigenomic features of normal myeloid cells and APL cells using integrative approaches including ChIA-PET for chromatin topology, ChIP-seq for epigenomic state, and RNA-seq for transcriptional output, to analyze the effects of the oncogenic fusion protein PML/RARα on the myeloid genomes. Significantly, we employed an inducible myeloid system, in which the expression of the PML/RARα protein is precisely control by ZnSO4 induction. With this system, we investigated the dynamic changes in chromatin topology triggered by nascent PML/RARα in the initial transformation stage, beginning at the normal myeloid state in leukemogenesis. We also analyzed the patient-derived APL cells harboring native PML/RARα to validate our observations in the inducible myeloid system.

Collectively, in this study we provided convincing data demonstrating that the PML/RARα proteins are aggressively involved in extensive chromatin interactions genome-wide in a specific manner. Although the DNA-binding properties of PML/RARα are derived from RARα, more than two thirds (2/3) of the PML/RARα binding loci did not overlap with RARα binding sites, indicating that this fusion protein acquired novel chromatin interacting capacities. Intriguingly, our data indicated that PML/RARα did not directly interfere with CTCF binding and chromatin looping, but rather that many of the PML/RARα-mediated chromatin loops overlapped the boundaries of CTCF-defined topological structures, and acted as a “stitch” or “staple” to interconnect separate chromatin topological structures into much larger domains with more condensed configuration, thereby reshaping the chromatin topology in normal myeloid cells leading to leukemogenesis.

Importantly, we also demonstrated that PML/RARα specifically intrude upon RNAPII-associated chromatin interaction domains of active genes in myeloid cells interrupting the binding of myeloid-specific TFs such as PU.1, IRF1, and CEBPB at enhancers and super-enhancers. The extensive chromatin binding and looping by PML/RARα could substantially compressed the chromatin topological structures around myeloid-specific transcriptional cassettes, thus leading to transcriptional repression of genes that are critical for myeloid differentiation and maturation. Perturbation experiments via induction (by ZnSO4) and depetion (by ATRA) of *in vivo* PML/RARα in PR9 cells to repress and to resecue the normal myeloid differentiation expression cassette further validated the specificity of PML/RARα-targeted genes. Additional perturbation experiments with native PML/RARα in patient-derived NB4 cells by ATRA treatment provided further evidence verifying PML/RARα’s target specificity in gene transcription reprression.

Taken together, our findings comprised a comprehensive view of the involvement of PML/RARα in chromatin topology during the early transformation process of PML/RARα-triggered APL genesis. Mechanistically, we posit that PML/RARα overrides the normal regulatory control of myeloid differentiation by reshaping the higher-order chromatin topology and compressing the transcriptional chromatin architectures. Therefore, the compressed chromatin domains would have reduced access by specific TFs and RNAPII, thus repress the transcription of genes critical to myeloid differentiation, and ultimately lead to leukemogensis. In sum, our results provide novel topological insights for the roles of PML/RARα in transforming myeloid cells into leukemia cells, likely a general mechanism for oncogenic fusion proteins in cancers.

## Methods

### Availability of data and materials

Genome-wide sequencing raw reads and processed files has been deposited at GEO. The accession number for the ChIA-PET, ChIP-Seq and RNA-Seq datasets for PR9 and NB4 cells reported in this paper is GEO: GSE137662. All datasets, materials and softwares used in this study are listed in the supplementary Table S1, S2 and S3, respectively.

### Cell Lines and Culture Conditions

PR9 (U937-PR9) cell line is a PML/RARα-inducible model constructed from U937, a normal myeloid precursor cell line without the t(15;17) translocation but expressing many myeloid-specific transcription factors important in myeloid development, including PU.1. To avoid the potential bias of clonal variations in culture, a single-cell subclone was selected. NB4 is an patient-derived APL cell line, carrying the t(15;17) translocation and expressing the PML/RARα fusion protein. Both PR9 and NB4 cells were cultured in RPMI 1640 (ThermoFisher, A10491), supplemented with 10% fetal bovine serum (ThermoFisher, 10082147). These cells were cultured at 37 °C, 5% CO2, and ambient oxygen levels.

ZnSO_4_ (Sigma, Z0251) was dissolved in sterile water as a stock solution at 100 mM. Induction for PML/RARα by ZnSO4 in PR9 cells: 100 μM ZnSO_4_ for 4 hours.

ATRA (Sigma, R2625) was dissolved in ethanol as a stock solution at 1 mM. ATRA treatment in NB4 cells: 10^−6^M ATRA for 24 or 48 hours.

### ChIA-PET library preparation

ChIA-PET libraries with antibody against PML, RARα, RNAPII, and CTCF were constructed using about 10^8^ input cells from PR9, PR9+Zn, and NB4 cell cultures, following the ChIA-PET protocol [29,30]. The ChIA-PET libraries were sequenced by paired-end reads using Illumina instrument.

### ChIP-Seq

In this study, we generated ChIP-seq data from PR9 and PR9+Zn cells for TFs of P300, PU.1, CEBPB, and IRF1, using the antibodies: anti-P300 (Abcam, ab14984), anti-PU.1 (Santa Cruz, sc-352X), anti-IRF1 antibody (Santa Cruz, sc-497x), anti-CEBPB antibody (Santa Cruz, sc-150x), and followed standard ChIP-seq protocol [8].

### RNA isolation and RNA-Seq library preparation

Total RNA was extracted with RNeasy mini kit (Qiagen, 74106) from the following cells: PR9 cells, PR9 cells treated with ZnSO4 (final concentration is 100 μM) at different time points (4h, 6h, 12h, 24h), PR9 cells pre-incubated with ZnSO4 for 4 hours and then treated with ATRA (final concentration is 1 μM) for another 24 or 48 hours, NB4 cells and NB4 cells treated with ATRA (final concentration is 1 μM) for 24 and 48 hours. Prior to RNA-Seq library preparations, rRNAs were depleted using Ribo-Zero rRNA removal kits (Illumina Inc, MRZH11124) from total RNA. Then, RNA libraries were prepared by ScriptSeq RNA-Seq library preparation kit (Illumina Inc, SSV21124). The RNA-Seq libraries were sequenced using NextSeq 500 platform for paired-end sequencing.

### 3D DNA FISH

The 3D DNA-FISH was performed with custom-synthesized oligonucleotides probes (MYcroarray) according to candidate genome loci (probe A: chr20: 31,261,904-31,361,904, probe B: chr20: 30,141,728-30,241,728) [16]. PR9 cells and PR9 cells treated with ZnSO4 (final concentration is 100 μM) for 4 hours were spin down onto a coverslip slide coating with poly-lysine for 20 minutes; then wash the slides with PBS for 3 times and air dried. The cells were fixed in methanol/acetic acid solution (3:1) for 5 minutes at 4 degrees, air dried, wash with PBS for 5 minutes. The cells were dehydration through an ethanol series (70%, 90%, 100%) and air-drying. Then the cells were permeabilized with 0.5% Triton X-100 in PBS on ice for 5 minutes, and wash with PBS for 5mintues. Customized FISH probe (MYcroarray) was warmed and mixed with hybridization buffer well. The cells and the probe mix were simultaneously subjected to DNA denaturation at 80°C for 5 min. The hybridization was performed at 37°C in the humid dark chamber for overnight. After coverslip removing, once washing of 10min at RT with 2×SSC/50% deionized formamide, pH 7.0, followed by once wash of 10 min at RT with 2×SSC and twice washes of 10 min at 55°C with 0.2×SSC were performed. Then cells on the slides were incubated with ProLong™ Gold Antifade Mountant with DAPI (ThermoFisher, 36931) in PBS buffer for 5 minutes and examined under the Leica SP8 confocal microscope. The distances between the probe pair were measure in 3D with IMARIS 9 software.

## Acknowledgements

The authors thank Dr. Oscar Junhong Luo and Dr. Guliang Li for initial data analysis, and Dr. Roel Verhaak for valuable comments on the manuscript.

## Author contributions

Y.R. conceptualized and supervised this study. P.W. designed the experiments and generated all genomic data with assistance from X.R. and M.Z. on library construction. Z.T., B.L., and S.Z.T. analyzed the data. J.J.Z and L.C. performed 3D DNA-FISH expriments and data analysis. P.S. and D.P. performed simulation and visualization of 3D chromatin folding models. P.W, Z.T, and Y.R. interpreted the results and wrote the manuscript with inputs from E.T.L and C-L.W.

## Corresponding author

Correspondence to Dr. Yijun Ruan.

## Ethics declarations

### Ethics approval and consent to participate

Not applicable

## Supplementary information

Supplemental information includes three tables and six figures can be found with this article.

