## supplemental Tables, figures and methods for "Chromatin Topology Reorganization and Transcription Repression by PML/RARα in Acute Promyeloid Leukemia"

Ping Wang^1^, Zhonghui Tang^1#^, Byoungkoo Lee^1^, Jacqueline Jufen Zhu^1,2^, Liuyang Cai^1^, Przemyslaw Szalaj^3^, Simon Zhongyuan Tian^1^, Meizhen Zheng^1^, Dariusz Plewczynski^3^, Xiaoan Ruan^1^, Edison T. Liu^1^, Chia-Lin Wei^1^, Yijun Ruan^1,2,*^

**SUPPLEMENTARY FIGURES** (Figure 1-6) …………………………………………………………………. P2

**SUPPLEMENTARY TABLES** (Table 1-3) ………………………………………………………………….. P10

**SUPPLEMENTARY METHODS** ……………………………………………………………………………... P13

**SUPPLEMENTARY FIGURES**

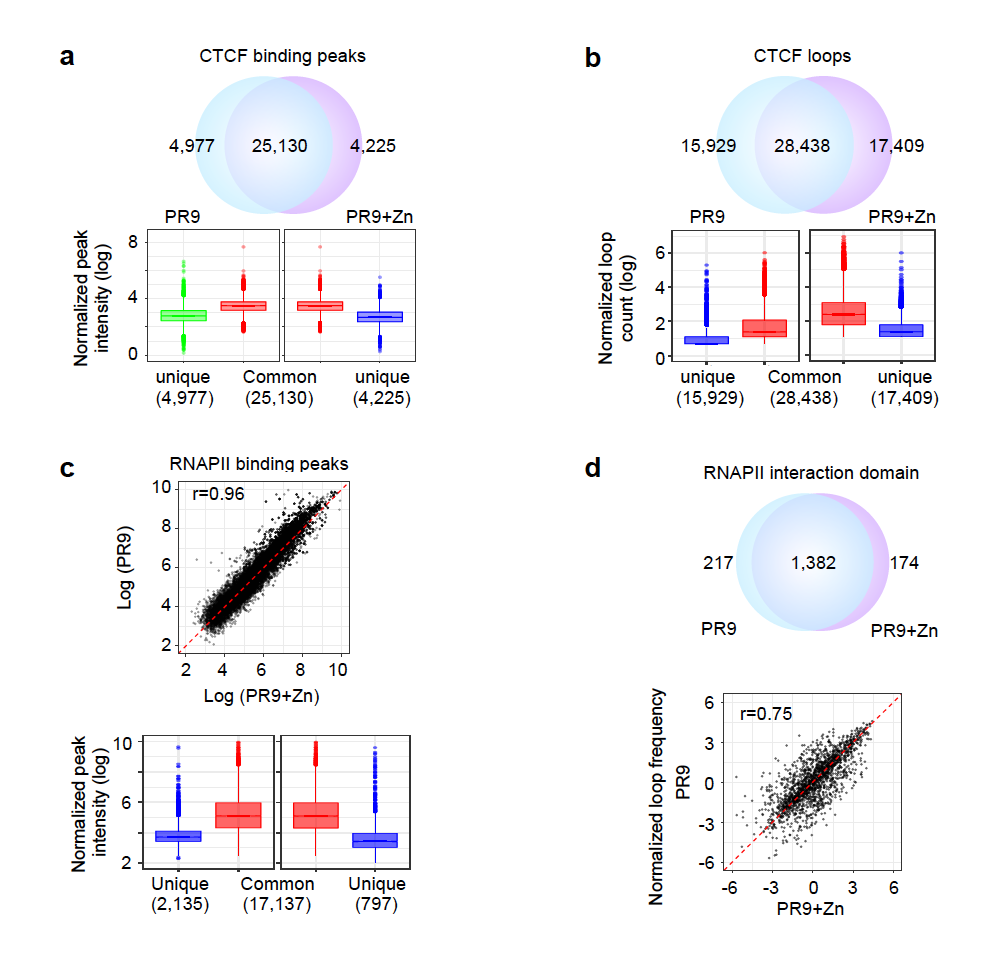

**Supplementary Figure 1**

**Supplementary Figure 1. Comparison of CTCF and RNAPII binding and looping in PR9 and PR9_Zn cells**. Related to Figure 1. **a-b.** CTCF binding peaks (a) and loops (b). Venn diagram shows common and unique CTCF peaks or loops between PR9 and PR9_Zn cells. Box plots show the intensities of peaks or loops that were unique or common to PR9 and PR9_Zn cells. **c.** Scatter plot of RNAPII binding peaks in PR9 and PR9_Zn cells (top). Box plots show the intensities of peaks that were unique or common to PR9 and PR9_Zn cells (bottom). **d.** Venn diagram shows RNAPII chromatin interaction domains between PR9 and PR9_Zn cells (top). Scatter plot displays the correlation of RNAPII looping frequency in PR9 and PR9_Zn cells (bottom).

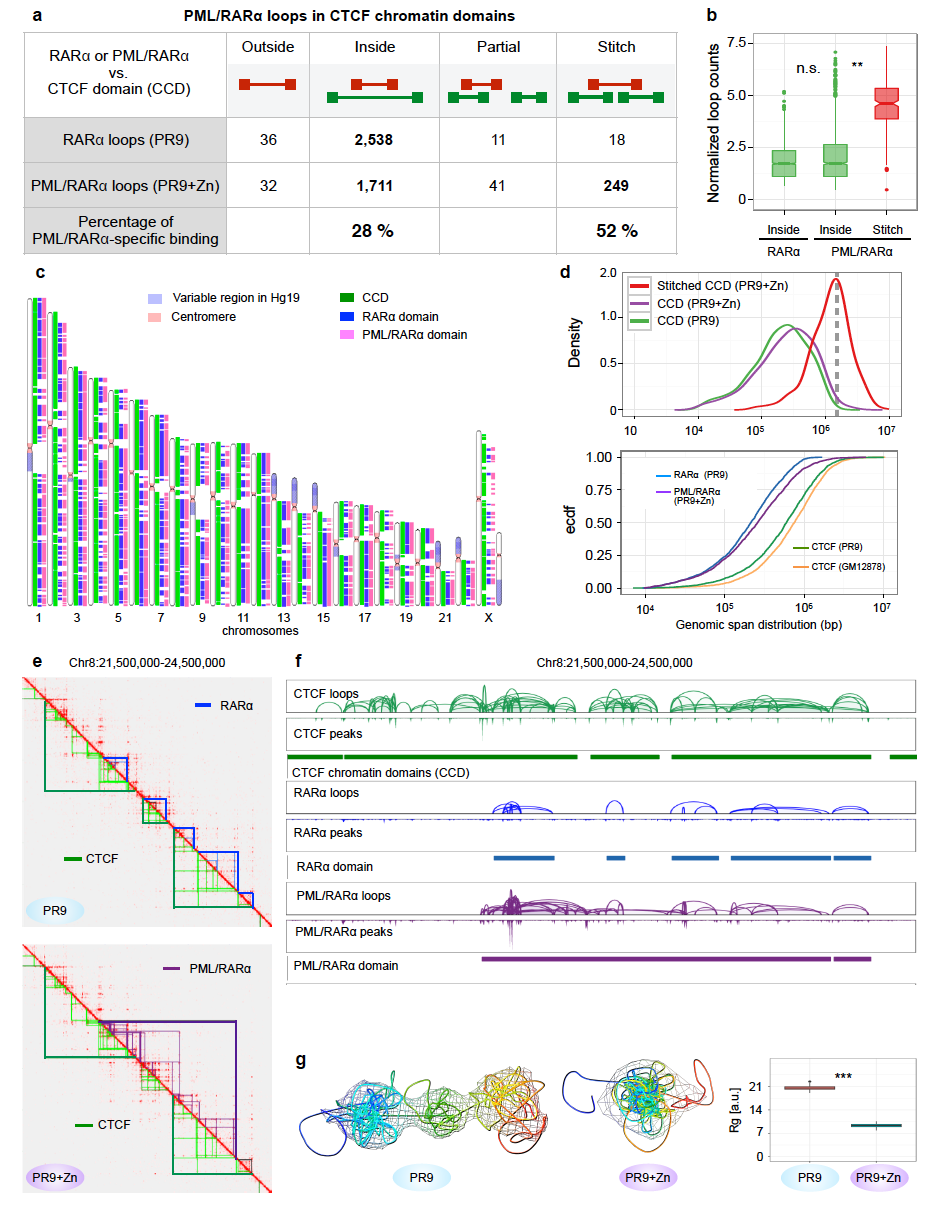

**Supplementary Figure 2**

**Supplementary Figure 2. PML/RARα-mediated chromatin interaction domain and impacts on the myeloid genome.** Related to Figure 2. **a.** Categorization of RARα in PR9 and PML/RARα in PR9+Zn cells in relation to CTCF-defined chromatin domains. **b.** Boxplot shows normalized chromatin contact frequency for RARα inside CCD domain in PR9 cells, and for PML/RARα inside CCD domain, as well as CCD domains stitched together in PR9+Zn cells. Kolmogorov-Smirnov test (K-S test) was used to test difference. ** for p < 0.001; n.s. for not significant (p > 0.001). **c.** Whole-genome view shows the distribution of CCD, RARα and PML/RARα domains in each chromosome. **d.** Size distribution CCD and “stitched” CCD in PR9 and PR9_Zn cells (top). The ECDF plot of chromatin loops for RARα, PML/RARα, and CTCF in PR9 and PR9_Zn cells, with the CTCF loops in GM12878 as a reference (bottom). **e.** Integrated 2D chromatin contact maps for a 3-Mb genomic segment from the combined (red) CTCF, RNAPII, and RARα ChIA-PET data in PR9 cells, as well as CTCF, RNAPII, and PML/RARα ChIA-PET data in PR9+Zn cells. Light green and dark green triangles indicate CTCF loop and CCD, respectively; light blue and dark blue triangles indicate RARα loops and domains in PR9 cells; and the light and dark purple triangles depict PML/RARα loops and domains in PR9 cells. **f.** Browser views of chromatin interaction loop, peaks, and chromatin domain of CTCF (green), RARα (blue), and PML/RARα (purple). **g.** 3D chromatin folding modeling for the genomic segment (same as in E and F): simulated average structure and ensemble in PR9 (left) and in PR9_Zn cells (middle). The boxplot shows radial diameter distributions of simulated 3D structures from 300 nuclei examined in PR9 and PR9_Zn cells. K-S test was used to test differences. ** *p* < 2.2e-16.

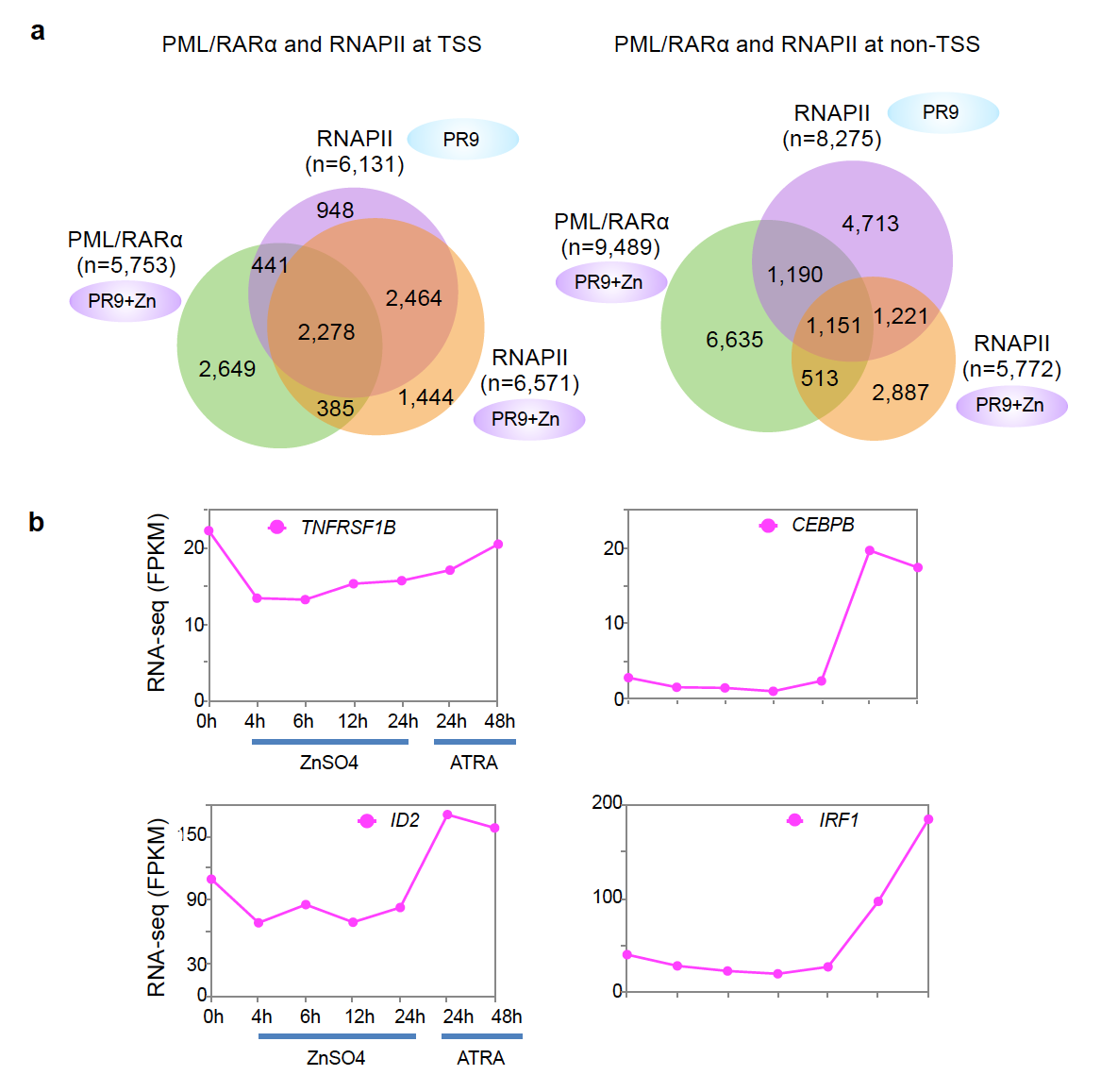

**Supplementary Figure 3**

**Supplementary Figure 3. Transcription repression of myeloid specific genes by PML/RARα.** Related to Figure 3. **a.** Venn diagrams show the overlap of binding sites by PML/RARα and RNAPII proximal to TSS of genes (left) and distal to TSS located at non-TSS loci (right). **b.** Line plots show gene (*IRF1*, *CEBPB*, *ID2*, *TNFRSF1B*) expression changes over the time course of ZnSO4 induction for PML/RARα activation and ATRA treatments in in PR9 cells.

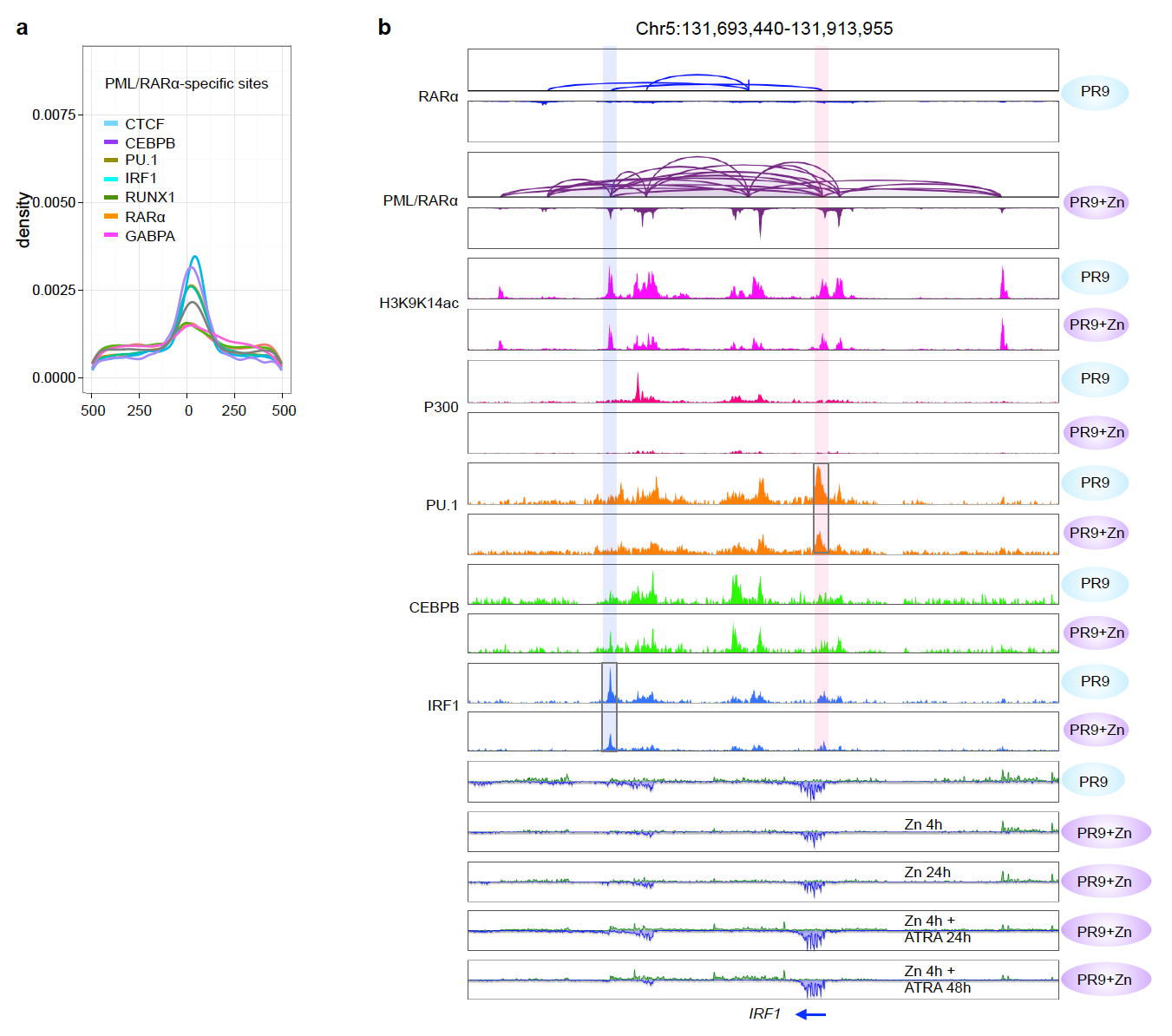

**Supplementary Figure 4**

**Supplementary Figure 4. Further details of motif analysis around PML/RARα binding sites.** Related to Figure 4. **a.** DNA binding motifs enriched (CTCF, CEBPB, PU.1, IRF1, RUNX1, RARA, and GABPA) at PML/RARα binding sites. **b.** An example browser view of chromatin interaction loops and binding peaks for RARα and PML/RARα,from ChIA-PET data, and the peaks from ChIP-seq data of H3K9K14ac and specific TFs at *IRF1* loci in both PR9 and PR9+Zn cells.

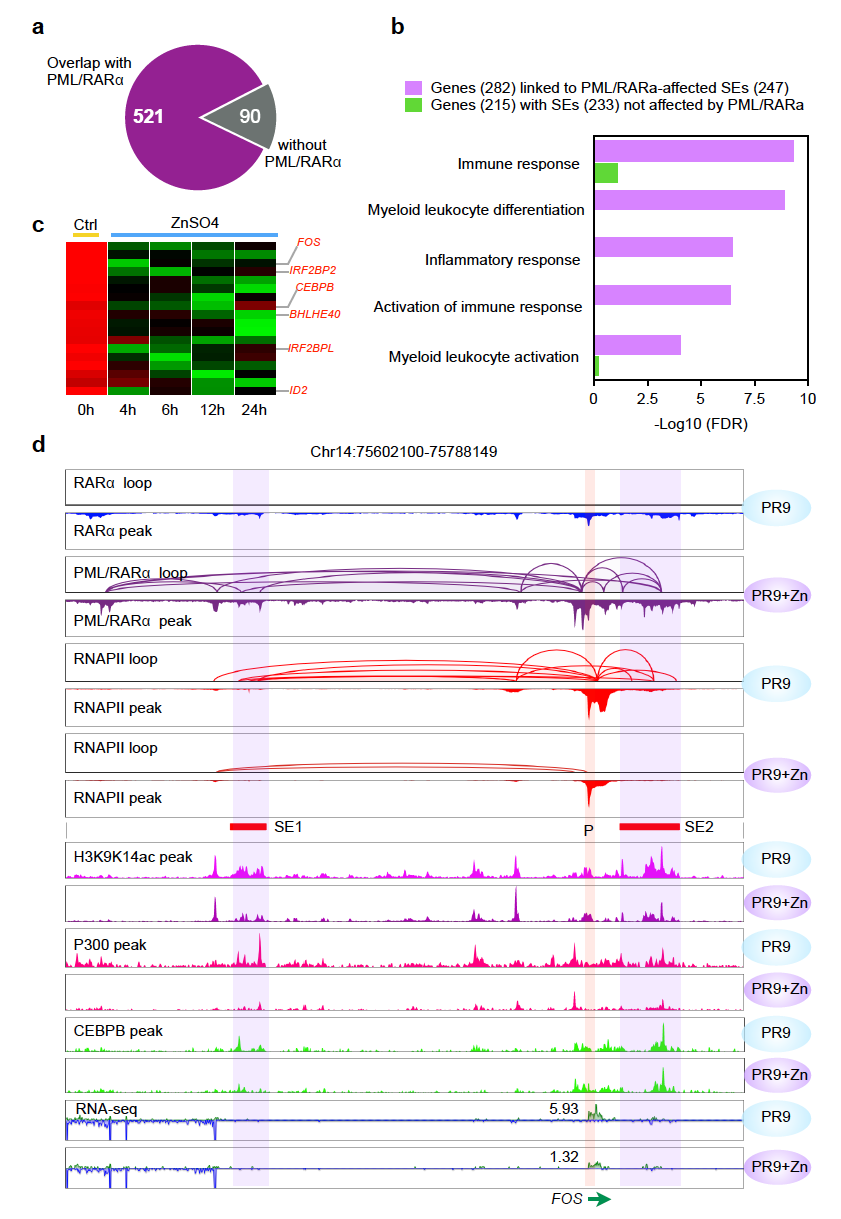

**Supplementary Figure 5**

**Supplementary Figure 5. Super-enhancers of myeloid specific genes affected by PML/RARα.** Related to Figure 5. **a.** Pie chart of SEs overlapped with PML/RARα anchors. **b.** GO enrichment analysis of genes (n=247) associated with SEs that were affected by PML/RARα, and for the genes associated with SEs that were not affected by PML/RARα. **c.** Heatmap shows expression profiles of PML/RARα-affected genes that are critical for myeloid development. **d.** An example of two SEs at their target gene *FOS* locus with tracks of PML/RARα and RNAPII ChIA-PET data (loops and peaks), along with ChIP-seq data of H3K9K14ac, P300 and CEBPB, as well as RNA-seq data from both PR9 and PR9+Zn cells. The locations of the promoter and the two SEs are highlighted.

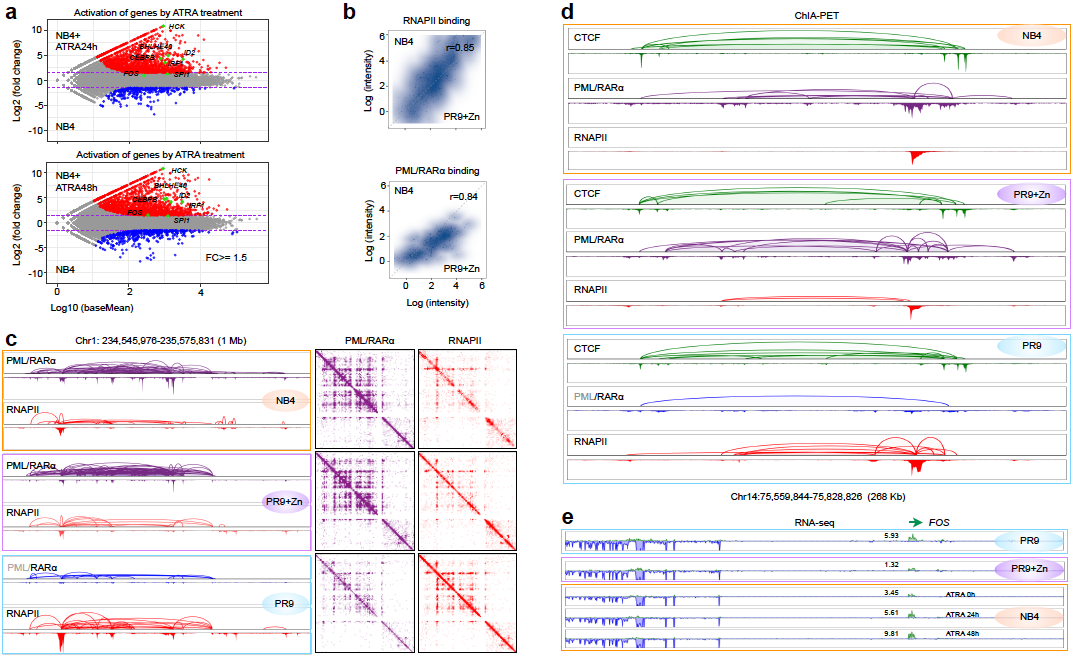

**Supplementary Figure 6**

**Supplementary Figure 6. Additional examples of PML/RARα in NB4 cell.** Related to Figure 6. **a.** Scatter plots show differentially expressed genes in NB4 cells and NB4 cells with ATRA treatment for 24 hours (top) and 48 hours (bottom). Red dots highlight genes activated by ATRA treatment, with representative genes HCK, BHLHE40, CEBPB, IRF1, FOS, and SPI1. **b.** Contour plots for correlations of RNAPII binding (up) and PML/RARα binding (low) intensity at the 119 target genes between NB4 vs. PR9+Zn cells. Correlation coefficient r-values are provided. **c.** Example shows comparable chromatin interaction profiles by PML/RARα and RNAPII in NB4 cells and PR9+Zn cells, whereas in PR9 cells, the RNAPII interaction data were much stronger. Left, browser view of loops and peaks. Right, 2D contact maps. **d.** Another example shows the local chromatin loops and peaks by CTCF (green), PML/RARα (purple) or RARα (blue), and RNAPII (red) at the *FOS* locus in NB4 cells (orange box), PR9+Zn cells (purple box), and PR9 cells (in blue box). **e.** *FOS* gene expression in PR9 and PR9+Zn cells, and in NB4 cells with ATRA treatment for 24 and 48 hours.

**SUPPLEMENTARY TABLES**

**Supplementary Table 1: Dataset used in this study**

| **ChIA-PET** | | | | | | |
| --- | --- | --- | --- | --- | --- | --- |
| Cell | Factor | Treatment | GEO accession | Uniquely mapped PETs | Intra-chr. PETs | Data source |
| PR9 | PML | control | GSE137662 | 10,376,163 | 616,023 | This study |
|  |  | ZnSO4 4h | GSE137662 | 61,255,598 | 5,757,440 |  |
|  | RAR𝛼 | control | GSE137662 | 15,608,754 | 3,084,399 |  |
|  |  | ZnSO4 4h | GSE137662 | 58,597,937 | 5,532,885 |  |
|  | RNAPII | control | GSE137662 | 180,539,278 | 11,194,276 |  |
|  |  | ZnSO4 4h | GSE137662 | 66,568,749 | 65,521,063 |  |
|  | CTCF | control | GSE137662 | 91,515,096 | 6,184,381 |  |
|  |  | ZnSO4 4h | GSE137662 | 36,355,723 | 4,255,073 |  |
| NB4 | PML | NA | GSE137662 | 97,332,648 | 34,880 |  |
|  | RAR𝛼 | NA | GSE137662 | 172,674,019 | 20,169 |  |
|  | RNAPII | NA | GSE137662 | 29,173,744 | 29,832 |  |
|  | CTCF | NA | GSE137662 | 22,674,016 | 58,471 |  |
| **ChIP-Seq** | | | | | | |
| Cell | Factor | Treatment | GEO accession | Uniquely mapped reads | Peak numbers (MACS) | Data source |
| PR9 | PU.1 | control | GSE137662 | 42,939,412 | 7,615 | This study |
|  |  | ZnSO4 4h | GSE137662 | 39,768,820 | 4,251 |  |
|  | CEBPB | control | GSE137662 | 43,253,760 | 3,879 |  |
|  |  | ZnSO4 4h | GSE137662 | 41,530,777 | 2,865 |  |
|  | IRF1 | control | GSE137662 | 45,188,813 | 3,829 |  |
|  |  | ZnSO4 h | GSE137662 | 47,838,743 | 2,456 |  |
|  | P300 | control | GSE137662 | 15,813,190 | 31,622 |  |
|  |  | ZnSO4 4h | GSE137662 | 9,150,658 | 16,479 |  |
|  | H3K9K14ac | control | GSM468215 | 11,261,531 | 30,150 | Martens *et al* ., 2010 |
|  |  | ZnSO4 4h | GSM468216 | 11,251,479 | 32,824 | Martens *et al* ., 2010 |
| **RNA-Seq (ribosomal RNA depleted)** | | | | | | |
| Cell | Experiment | Treatment | GEO accession | Uniquely mapped reads |  | Data source |
| PR9 |  | control | GSE137662 | 108,254,542 |  | This study |
|  |  | ZnSO4 4h | GSE137662 | 108,122,444 |  |  |
|  |  | ZnSO4 6h | GSE137662 | 112,468,120 |  |  |
|  |  | ZnSO4 12h | GSE137662 | 89,297,626 |  |  |
|  |  | ZnSO4 24h | GSE137662 | 107,669,338 |  |  |
|  |  | ZnSO4 4h+ATRA 24h | GSE137662 | 63,868,024 |  |  |
|  |  | ZnSO4 4h+ATRA 48h | GSE137662 | 50,140,652 |  |  |
| NB4 |  | control | GSE137662 | 59,455,644 |  |  |
|  |  | ATRA 24h | GSE137662 | 57,918,501 |  |  |
|  |  | ATRA 48h | GSE137662 | 59,042,779 |  |  |

**Supplementary Table 2: Reagents used in this study**

| Resource | Source | Identifier | Experiments or analysis |
| --- | --- | --- | --- |
| PML antibody | Santa Cruz | Cat# sc-966X | ChIA-PET |
| RARα antibody | Santa Cruz | Cat# sc-551X | ChIA-PET |
| RNA Polymerase II antibody | Biolegend | Cat# 664912 | ChIA-PET |
| CTCF antibody | Abcam | Cat# ab70303 | ChIA-PET |
| P300 antibody | Abcam | Cat# ab14984 | ChIP-Seq |
| PU.1 antibody | Santa Cruz | Cat# sc-352X | ChIP-Seq |
| CEBPB antibody | Santa Cruz | Cat# sc-150X | ChIP-Seq |
| IRF1 antibody | Santa Cruz | Cat# sc-497X | ChIP-Seq |
| Nextera DNA sample preparation kit | Illumina, Inc | Cat# FC-121-1030 | ChIA-PET |
| Nextera Index kit | Illumina, Inc | Cat# FC-121-1011 | ChIA-PET |
| ScriptSeq RNA-seq library preparation kit | Illumina, Inc | Cat# SSV21124 | RNA-seq |
| Ribo-Zero rRNA Removal Kit (Human/Mouse/Rat) | Illumina, Inc | Cat# MRZH11124 | RNA-seq |
| TruSeq ChIP library preparation kit -Set A | Illumina, Inc | Cat# IP-202-1012 | ChIP-Seq |
| LightCycler 480 SYBR Green I Master | Roche | Cat# 04887352001 | ChIP-Seq, ChIA-PET |
| DNA Clean & Concentrator-5 | Zymo research | Cat# 4013 | ChIA-PET |
| QIAprep Spin Miniprep Kit | Qiagen | Cat# 27106 | ChIA-PET, ChIP-seq |
| RNeasy Mini Kit | Qiagen | Cat# 74106 | RNA-seq |
| ZnSO4⋅7H2O | Sigma | Cat# Z0251 | Cell culture |
| All-trans retinoic acid (ATRA) | Sigma | Cat# R2625 | Cell culture |
| EGS (ethylene glycol bis(succinimidyl succinate)) | ThermoFisher | Cat# 21565 | ChIA-PET |
| Formaldehyde | Millipore | Cat# 104003 | ChIA-PET, ChIP-seq |
| Customized FISH probe | MYcroarray | NA | 3D-DNA FISH |
| T4 DNA polymerase | Promega | Cat# 4421 | ChIA-PET |
| Klenow Fragment (3’-5’ Exo) | NEB | Cat# M0212M | ChIA-PET |
| T4 DNA Ligase, HC (30 U/µL) | ThermoFisher | Cat# EL0013 | ChIA-PET |
| T4 DNA Ligase Buffer | ThermoFisher | Cat# 46300018 | ChIA-PET |
| T4 DNA Ligase Reaction Buffer | NEB | Cat# B0202S | ChIA-PET |
| DNA Polymerase I (*E. coli*) | NEB | Cat# M0209L | ChIA-PET |
| Dynabeads™ Protein G | ThermoFisher | Cat# 10009D | ChIA-PET |
| AMPURE XP beads | Beckman Coulter | Cat# A63881 | ChIA-PET |
| Dynabeads™ M-280 Streptavidin | ThermoFisher | Cat# 11206D | ChIA-PET, ChIP-seq |
| 2% Agarose Gel Cassette | Sage Science | Cat# BDF2010 | ChIA-PET |
| ProLong™ Gold Antifade Mountant with DAPI | ThermoFisher | Cat# P36931 | 3D-DNA FISH |
| MmeI | NEB | Cat# R0637l | ChIA-PET |
| Proteinase K | ThermoFisher | Cat# 25530049 | ChIA-PET, ChIP-Seq |
| S-adenosylmethionine (SAM) | NEB | Cat# B9003S | ChIA-PET |
| GlycoBlue™ Coprecipitant | ThermoFisher | Cat# AM9516 | ChIA-PET |
| Sodium Acetate (3 M), pH 5.5, RNase-free | ThermoFisher | Cat# AM9740 | ChIA-PET |
| 2% Agarose Gel Cassettes | Sage Science | Cat# BDF2010 | ChIA-PET |

**Supplementary Table 3: Software and Algorithms used in this study**

| Resource (Reference) | Identifier |
| --- | --- |
| Bowtie2[1] | <http://bowtie-bio.sourceforge.net/index.shtml> |
| Picard (Broad Institute) | <http://broadinstitute.github.io/picard/> |
| Edge R[2] | https://bioconductor.org/packages/release/bioc/html/edgeR.html |
| MACS[3] | <https://github.com/taoliu/MACS> |
| 3D-GNOME[4] | <http://nucleus3d.cent.uw.edu.pl/> |
| GREAT[5] | <http://great.stanford.edu/public/html/> |
| IMARIS 9 (BITPLANE) | http://www.bitplane.com/releasenotes/imaris930.aspx |
| Juicer/Juicebox[6,7] | <http://aidenlab.org/juicebox/> |
| JASPAR[8] | <http://jaspar.genereg.net> |
| ROSE[9,10] | [http://younglab.wi.mit](http://younglab.wi.mit.edu/super%20_enhancer_code.html).edu/super_enhancer_code.html |

**SUPPLEMENTAYT METHODS**

**ChIA-PET data processing and clustering of interaction PETs**

In this study, ChIA-PET data was produced by the long-reads ChIA-PET and short-reads ChIA-PET protocols. The long-read ChIA-PET and short-read ChIA-PET data sets were processed using an updated ChIA-PET data processing pipeline, called ChIA-PIPE [11]. From each library data, protein binding peaks, chromatin loops and chromatin interaction domains were identified.

**Comparison of ChIA-PET data for calling of binding peaks, loops, and chromatin domains**

To compare the CTCF ChIA-PET data, we intersected the CTCF peak regions between the PR9 and PR9+Zn data, and identify unique and common sets of the peaks (Supplementary Fig. 1d). For each condition, we sampled the same number of PET to calculate CTCF binding intensity at each peak region (Supplementary Fig. 1a). To compare the loops, we extended ±1.5 kb to each anchor of CTCF loops to intersect common loops. Loop showing both overlapped anchors with that of a loop from different datasets was considered as a common loop (Supplementary Fig. 1b).

Considering that the CTCF ChIA-PET data showed the high similarity between PR9 and PR9+Zn cells, we merged the two datasets together, and used the combined data to generate a reference 3D genome map mediated by CTCF for myeloid cell PR9, which includes CTCF peaks, CTCF loops, CTCF chromatin domains (CCD) similar as we did for the 3D genome map of GM12878 (Tang et al., 2015). The same approach was also applied to characterize RARα, RNAPII and PML ChIA-PET data generated from PR9 and PR9+Zn cells. The results were shown in Fig. 2a.

**Identification and classification of RARα and PML/RARα binding sites**

We generated PML and RARα ChIA-PET libraries for PR9 and PR9+Zn cells, respectively. Using ChIA-PIPE, we called peaks and loops for corresponding factors. As expected, no significant signals for peaks and loops were detected PML ChIA-PET data from PR9 cells, because PML is not a chromatin protein. The binding peaks detected in the RARα ChIA-PET data in PR9 represented the endogenous RARα binding sites. In PR9+Zn cells, the binding peaks detected by RARα ChIA-PET data could for both of the endogenous RARα and the induced PML/RARα protein, while the binding peaks identified by PML ChIA-PET data should only represent the induced PML/RARα fusion protein. To identify high-confidence data, we considered only the peak regions detected more than two datasets in these three ChIA-PET experiments in PR9 and PR9+Zn cells as illustrated in Fig. 2b. According to binding patterns, the peak regions were classified into three categories (Fig. 2b): endogenous RARα-specific peaks (Type I), shared peaks by endogenous RARα and the induced PML/RARα (Type II), and PML/RARα-specific peaks (Type III). In type II category, the endogenous RARα and the induced PML/RARα were co-occupied at the same genomic regions.

**Identification and characterization of PML/RARα-mediated chromatin interactions**

The PML- and RARα- ChIA-PET datasets essentially represent the detection of the fusion protein PML/RARα. To comprehensively analyze the chromatin interactions mediated by PML/RARα, we merged two datasets together. The combined PETs were used to identify high-confidence PML/RARα loops and domain similar as we did for CTCF (Fig. 2a).

**Visualization of 2D contact map and BASIC browser**

The pairwise contact data from short-read ChIA-PET and long-read ChIA-PET were combined. We produced 2D contact map files (hic files) using Juicer tools [7] (version 1.6.1) (<https://github.com/aidenlab/juicer/wiki/Download>), and visualized them using Juicebox (version 1.6.1) [6]. Ten different resolutions were applied from 2.5Mb to 10Kb. We chose none normalization in Juicebox for our ChIA-PET 2D contact map visualization. We generated contact map files for individual single-factor conditions, CTCF, RNAPII, PML/RARα, and RARα for control and treatment cases, respectively. In ChIA-PIPE, there is a BASIC browser viewer for visualizing loops and peaks for each ChIA-PET dataset.

**3D structure modeling, measurement of 3D structure condensation and visualization**

The modeling was performed using 3D-GNOME software we developed earlier [4]. The models were generated using a combined CTCF, RNAPII, and PML/RARα ChIA-PET loop dataset for control and treatment cases, respectively. The PET count threshold was set to 3 for CTCF and PML/RARα loops and 2 for RNAPII loops. Loops, which PET count is less than the threshold, were filtered out. Also, we took the loops, where both loop anchors are inside of the target genomic region, as an input for 3D-GNOME. Neither anchor or sub anchor heatmaps nor CTCF motifs orientations optional energy terms were used in the modeling. An ensemble of 100 structures was created for each target genomic region. The 3D polymer models from 3D-GNOME are based on chromatin loops, and thus the genomic distances between consecutive beads in each model are variable. To assure uniform bead coverage over the target genomic region, the 3D model structures were interpolated using a cubic spline method with a 1Kb interval. 3D visualizations (Fig. 2h and Supplementary Fig. 2g) were created using the UCSF Chimera package. In every panel, the medoid structure (i.e. a structure for which the average pairwise dissimilarity to all the other structures is minimal; medoid can be intuitively thought of as a central or average structure) is presented, with the color corresponding to the genomic position. Additionally, an ensemble cloud that represents the average conformation of the whole ensemble of structures is shown. The cloud was calculated using the Calculate Occupancy tool in Chimera (the structures were previously co-aligned). While the locations of the individual loops varied between individual models, the ensemble clouds demonstrate that the overall shapes of 100 model structures were homogenous. Generally, the control structures consist of multiple, well-separated domains connected by linker regions, whereas the treatment structures are compacted to a single globule. To compare the level of compaction between control and treatment structures, we calculated the radius of gyration for individual structures (Fig. 2h and Supplementary Fig. 2g).

**Identification of binding motifs at PML/RARα peak regions**

The DNA sequences in PML/RARα binding peaks were extracted for scanning protein binding consensus sequences. The position weight matrix for RARα, PU.1, IRF1, CEBPB, RUNX1, GABPA and CTCF motifs were downloaded from the JASPAR database (<http://jaspar.genereg.net>) [8]. FIMO from MEME suite (version 4.12) was applied to identify of consensus sequences for each protein factor. The threshold of p-value 10^-4^ was applied to filter output of FIMO. The occurrence frequency of consensus sequence for each protein factor was plotted surrounding RARα consensus sequence in a 1kb window (Fig. 4a and Supplementary Fig. 4a). The plot shows consensus sequences of transcription factors, including PU.1, IRF1 and CEBPB, are proximal to the consensus sequences of RARα in PML/RARα binding regions, which suggests PML/RARα may occupy same regions with transcription factors.

**Comparison of RNAPII-mediated chromatin interactions between control and treatment condition**

We described the procedures of identifying RNAPII chromatin interaction domains with significant changes in RNAPII-mediated chromatin interactions. We first called the union regions of joint RNAPII chromatin interaction domains from control and treatment. We then calculated numbers of inter-ligation PETs in the union regions at control and treatment, respectively. Similarly, the accumulated PET counts of PET clusters in the union regions were also calculated at control and treatment, respectively. We treated control and treatment conditions as the two groups, and inter-ligation PETs and clustered PETs as the two categories, generating a 2 X 2 contingency table as shown below for Chi-Square test. The R package chisq.test was applied to calculate the p-value. The RNAPII chromatin contact domain was considered with a significant change in chromatin interaction when p-value less than 0.01.

| **Group** | **Categories** | |
| --- | --- | --- |
|  | **inter-ligation PETs** | **clustered PETs from loops** |
| Control | N_a_ | N_b_ |
| Treatment | M_a_ | M_b_ |

Statistical numbers of RNAPII involved loop difference between PR9 cells and ZnSO4 treated PR9 cells at the local regions were summarized in Supplementary Fig. 4b, and *p*-values were estimated from two-side Chi-square test.

**RNA-Seq and differential gene expression analysis**

Sequenced reads were mapped to the human reference genome hg19 using Bowtie2 (<http://bowtie-bio.sourceforge.net/bowtie2/index.shtml>) [1] with default parameters. Only uniquely mapped reads were detained for the downstream analysis. The counts of mapped reads fell in exons of each gene were calculated and were then applied to identify differentially expressed genes using R package ‘edge R’ [2]. The expression values for each gene was normalized using reads per kilobase per million (FPKM) mapped reads. The expression heat maps were generated with FPKM values of genes using R package ‘heatmap2’ (Fig. 3b). The log fold-change of 1 and *p*-value of 0.01 were applied to identify differentially expressed gene between control and treatment conditions in PR9 cells. The gene ontology enrichment analysis was performed for differentially expressed genes using GREAT (<http://great.stanford.edu/public/html/>) (Fig. 3c). To compare the similarity in gene expression between PR9 and NB4 cells in a variety of conditions, genes with FPKM >= 1 in both PR9 and NB4 cells were applied to calculate Pearson correlation using R package stats (Fig. 6b).

**ChIP-Seq analysis**

Sequenced reads were mapped to the human reference genome hg19 using bwa-mem (version 0.7.16) with default parameters. Only uniquely mapped reads were detained for the downstream analysis. In order to compare the ChIP-Seq profiles between control and treatment, we sampled the same number of mapped reads for peak calling in both control and treatment conditions. MACS 1.4.2 was used for identification of peaks using default parameters. If genomic distance between peaks is less than 2 kb, the peaks were merged together and applied for downstream analysis. To compare the correlation of RNAPII and PML/RARα binding profiles between PR9 and NB4 cells, the values of reads per million (RPM) for peak regions were applied to calculate Pearson correlation using R package stats (Fig. 6c).

**Identification and characterization of super-enhancers**

We used ROSE (<http://younglab.wi.mit.edu/super_enhancer_code.html>) [9,10] with default setting to discovery super-enhancer based on genome-wide H3K9K14ac profile. Super-enhancers were identified as regions with slope denoting (focus ratio)/(region size annotation enhancer) greater than 1. The super-enhancer regions were identified in control and treatment condition, respectively (Fig. 5b). The super-enhancer regions overlapped more than 70% between conditions were categorized as common super-enhancers. Except the common super-enhancers, others were grouped as condition-specific super-enhancers. The categorized super-enhancer regions were intersected with PML/RARα binding peaks, and only super-enhancer regions overlapped with PML/RARα binding peaks were detained for further characterization (Supplementary Fig. 6a, Fig. 6c).

**Quantification and statistical analysis methods**

Kolmogorov-Smirnov test (K-S test) was applied in Figure 2h, Supplementary Figure 2b, Supplementary Figure 2g and Figure 5d. Mann-Whitney u test was applied in Figure 2g. All statistical tests used two-side tests.
